# Press Releases Shape Online Attention for Neuroscience Articles

**DOI:** 10.1101/2025.10.31.685745

**Authors:** R. Bastida-Barau, J. Giner-Miguelez, C. Barcia, S. Ferré

## Abstract

Alternative indexes (altmetrics), such as the number of social media posts or news mentions, assess the relevance of a publication beyond academia. At the same time, composite metrics, like Altmetric Attention Score (AAS), gather these indicators to provide an overall measure of the online attention of an article. Communication departments at research institutions aim to help publications reach the general public and, therefore, play a key role in improving the altmetrics performance of scientific articles. However, the most effective strategies to achieve this remain unclear. No standardized protocols exist to guide communication professionals in maximizing the impact of their outreach efforts. Thus, in the present research, carried out by the communication department of a neuroscience research center, we conducted three complementary analyses to help to fill this gap. First, a study involving more than 3,000 neuroscience publications from 8 similar research centers revealed correlations between altmetrics and citations. Second, a retrospective analysis of 201 in-house articles found that active communication campaigns were associated with better altmetric performance, with notable differences between publications that included a press release (PR) and those that did not. Finally, a prospective experiment involving 25 articles examined changes in altmetrics before and after the dissemination of a PR or posting on X. Results showed that PRs significantly increased the number of news mentions and X posts, leading to an average rise of approximately 70 points in the AAS. These findings highlight the importance of PRs in maximizing the societal visibility and impact of scientific publications.

## 1. INTRODUCTION

A major challenge in research evaluation is identifying new indicators that better capture the true impact of research outputs (European Commission, 2023). Bibliometric indexes, such as the impact factor (IF), percentile ranks, or the number of citations, have commonly been used to measure how a publication contributes to scientific knowledge. However, they are mostly restricted to the academic world. Also, because of their limitations (Adler et al., 2009; Hicks et al., 2015; Rossner et al., 2008; Seglen, 1997; Vanclay, 2012), public institutions and editorials have joint efforts to identify new tools to better evaluate the scientific and social impact of a publication (ASCB, 2012; CoARA, n.d.; Nature Editors, 2005; Plos Medicine Editors, 2006).

In the Declaration on Research Assessment (DORA), a group of editors and publishers of scientific journals claimed the need to improve how researchers and their outputs are evaluated. This document, created at the Annual Meeting of The American Society for Cell Biology in 2012, and currently signed by more than 22,000 individuals and organizations, highlights, among other points: 1) the need to assess research on its own merits (and not on the prestige of the journal where it is published), and 2) exploring other indicators of significance (ASCB, 2012).

In the context of rethinking how impact should be measured, new tools have emerged to provide additional perspectives for quantifying the interest in a publication. Whereas traditional bibliometric indexes neglect impact outside academia (Priem et al., 2010), alternative indexes (altmetrics), such as the number of X posts (formerly Twitter) or news mentions referring to an article, aim to instantly include online conversations happening around a publication among the general public, supplying qualitative data that complement traditional citation-based metrics. Also, composite altmetrics appeared, such as PlumX or the Altmetric Attention Score (AAS), integrating multiple of these indicators to provide a combined view of attention across different sources (Altmetric, n.d.; Altmetric, 2023; Elsevier, n.d.; Scarlat et al., 2015).

Scientific publishers are increasingly implementing these composite altmetrics tools on their online platforms. Currently, the AAS is available, among others, on ACS, Cambridge University Press, Springer Nature, Taylor & Francis and Wiley, while PlumX is on Elsevier. These platforms also often provide guidelines with outreach plans to achieve greater attention (e.g., Cambridge University Press, n.d.; Clark, 2025; MDPI, n.d.; Springer Nature, n.d.; Taylor & Francis Author Services, n.d.) and send recommendations to their authors on how to spread their articles using social media (SM). Nonetheless, it is not clear on what studies these recommendations are based. Research in this field is scarce; hence, scientists and communication professionals do not have much objective data available to optimize this task.

Apart from SM, news coverage in the press continues to be an important channel through which the general public accesses information on scientific advancements. However, what makes an article gain media attention remains a complex issue. A classic highly-cited study from 1998 found that most newspaper stories covered articles that had been issued a press release (PR) (de Semir et al., 1998), and more recent studies showed that PRs are linked to higher volume of newspaper coverage (Cho & Yoon, 2019; Stryker, 2002). In line with this, Fuoco et al. (2023) found in a retrospective study analysing environmental health research papers that the presence of a PR was strongly associated with higher AAS. Nonetheless, as Stryker (2002) points out, this may be due to the fact that the most newsworthy articles are more likely to have an associated PR, rather than the PR itself exerting a direct effect. Prospective experiments, such as the one presented in the third part of this article, which measure various parameters before and after a campaign, are needed to clarify the actual impact of a PR on the number of news mentions and AAS, as well as on SM attention, which has not been extensively investigated.

Another important but unresolved question is whether increased public interest (which may be fostered by an effective communication campaign) leads to a higher number of citations. While several studies have found correlations between public attention and citation counts (Araujo et al., 2021; Bardus et al., 2020; Costas et al., 2015; Eysenbach, 2011; Maggio et al., 2018; Knight, 2014; O’Connor et al., 2017; Patthi et al., 2017; Wang et al., 2024), others have not (Allen et al., 2013; Branch et al., 2024; Delli et al., 2017; Fox et al., 2015, 2016; Livas & Delli, 2018; Peters et al., 2016; Tonia et al., 2016). Further research may provide clearer insights into the complex interrelationships between social exposure, readership and/or downloads, and citation counts across different research fields.

In the present study, we first performed a retrospective analysis of more than 3,000 articles and reviews in the neuroscience field, looking for correlations between social attention and citations. Since it typically takes a few years for published articles to reach their peak citation count (Schloegl & Gorraiz, 2011), we only included articles that were at least four years old at the time of analysis.

In a second part, we examined the social impact of 201 articles from our center, the *Institut de Neurociències de la Universitat Autònoma de Barcelona (INc-UAB)*, categorized by the communication actions carried out by the outreach department. Our goal was to determine if an active communication campaign correlated with a higher online attention, measured with different altmetric indicators.

Finally, to establish causal relationships, a prospective experiment involving 25 articles generated in our institution was conducted, examining altmetrics variations before and after issuing a PR or posting a publication on platform X.

## 2. METHODS

In this section, we detail the sample selection process for the retrospective analysis, including the inclusion and exclusion criteria, as well as the approach taken for the prospective study. Lastly, we describe the statistical analysis conducted in each case.

### 2.1 Selection of publications from 8 worldwide Neuroscience Research Centers

For the first retrospective analysis, we selected seven national and international neuroscience research institutions based on similar size and a certain degree of geographical dispersion: the Achucarro Basque Center for Neuroscience (Spain), the Instituto de Neurociencias de Alicante (Spain), the Centro Internacional de Neurociencia Cajal (Spain), the Institute of Neuroscience of la Timone (France), the Neurocentre Magendie (France), the Neuroscience Research Australia (Australia), and the UCL Neuroscience (UK).

From a bibliographical search according to the affiliation (using Scopus in December 2021), we obtained all the articles and reviews published from 2017 to 2020 by the selected centers. We gathered 4,233 publications, to which we added our in-house articles and reviews from 2017 to 2020 (586 publications). The bibliographic search by Scopus provided the DOI, type of publication, title, authors, year, and source title for each item. The IF was obtained from the Journal of Citations Reports from Web of Science (WoS, Clarivate), considering the year when each article was published. Using the Altmetric Application Programming Interface (API) in December 2021, we collected the number of mentions in the news, number of X posts, number of Facebook posts, number of readers on Mendeley (which is the amount of Mendeley users that have added a particular document to a Mendeley library), AAS, and AAS percentile within the same journal. It is important to indicate that, according to Bar-Ilan and colleagues (Bar-Ilan et al., 2019), Altmetric exclusively monitors Mendeley readership for research outputs that have already been referenced on at least one social network, disregarding a portion of the audience. We did not analyze Reddit, Blogs, F1000Prime, Video, Wikipedia, Peer review, and Policy documents, because their impact is residual. However, as they are included in the AAS, they indirectly have been studied. We did not cover Instagram (IG), Bluesky, LinkedIn, Google+, Pinterest, CiteULike, or Sina Weibo, because Altmetric was not tracking these platforms at the moment the study was performed. Using the PlumX API, we obtained the number of citations, abstract views, tweets, and Facebook posts. However, some data from the PlumX API did not match what we observed on X or Facebook, and we discarded the source. Following this criteria and workflow (**Fig. 1**), and keeping only those articles and reviews detected by the APIs, we finally collected 3,129 publications (2,745 articles and 384 reviews). Finally, in March 2025, we obtained from Scopus the number of citations for each article.

**Fig. 1.**
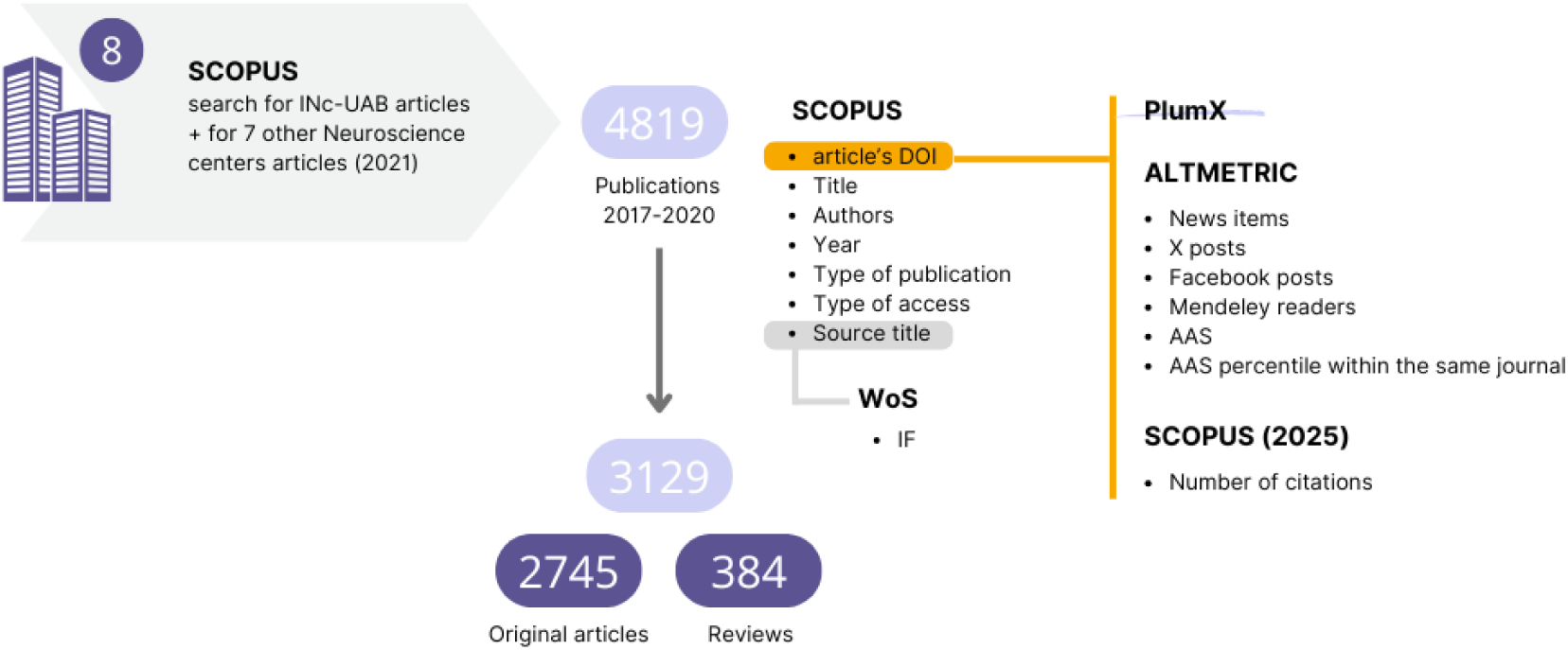
General Procedure for Data Extraction of Publications from Eight International Neuroscience Research Centers. We conducted a comprehensive search on Scopus, targeting the affiliations of seven neuroscience research institutions, which yielded a total of 4,233 publications from 2017 to 2020. In addition, we incorporated 586 in-house articles and reviews published during the same timeframe. A first Scopus search provided essential details for each publication, including the DOI, publication type, title, authors, year, and source title. The IF was obtained from the Web of Science (Clarivate). The Altmetric API supplied the metrics such as mentions in the news, posts on X, Facebook engagements, Mendeley readers, AAS, and AAS percentile within the corresponding journal for each article. Data retrieved from the PlumX API was discarded. After excluding publications not identified by both the Altmetric and PlumX APIs, we finalized our dataset with 3,129 publications, consisting of 2,745 articles and 384 reviews. From a second search in Scopus in March 2025 we obtained the number of citations, at least four years after the publication of the articles.

### 2.2 Selection of in-house publications

For the second analysis, a Scopus search by author ID, of 61 INc-UAB principal investigators, was performed in October 2021, obtaining a list of 718 publications from 2016 to 2020. To increase the sample, a second search was done in January 2023, obtaining 182 additional publications from 2021. We only selected original articles in English where an INc-UAB member was the principal author (only first or last author contributions were considered, according to the criteria of leadership in the neuroscience field), resulting in 201 publications.

As all principal authors were in-house members, our press department was in charge of the communication campaign for each article, and the outreach strategy followed in each case was controlled and perfectly known. Hence, we divided the publications into two groups: those that were subject of a PR (PR group, n=28) and those that were not (NPR group, n=173).

We obtained the DOI, title, authors, year, source title, and citation count from Scopus (January 2023). The IF was obtained from the Journal of Citations Reports from Web of Science (WoS, Clarivate), considering the year when each article was published. Using the Altmetric API (Dec. 2021 and Jan. 23), we collected the number of mentions in the news, X posts, Facebook posts, Mendeley readers, AAS, and AAS percentile within the same journal. Finally, in March 2025, we obtained from Scopus the number of citations for each article (**Fig. 2**).

**Fig. 2.**
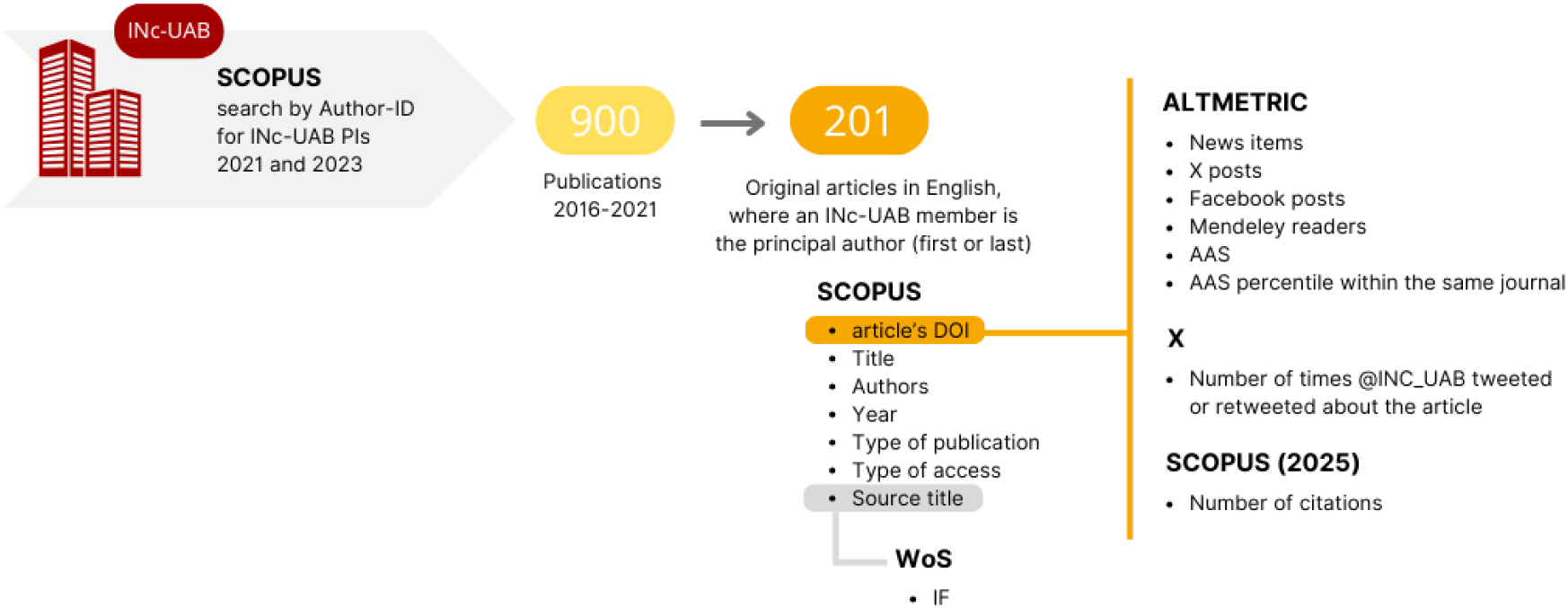
General Procedure for Data Extraction of Publications by INc-UAB Members (2016-2021). We performed two searches on Scopus using the author IDs of 61 principal investigators affiliated to INc-UAB: one in October 2021 and another in January 2023. This yielded a total of 900 publications from 2016 to 2021. We retained only original articles published in English, with an INc-UAB member as the principal author, resulting in a final dataset of 201 articles. Key details, including the DOI, title, authors, year, source title, type of access, and citation count, were extracted from Scopus. The Impact Factor was sourced from the Web of Science (WoS). Additionally, using the Altmetric API, we collected metrics such as mentions in the news, posts on X, Facebook interactions, Mendeley readers, AAS, and AAS percentile within the respective journal. X API provided with the tweets and retweets mentioning each article. Finally, from a second search in Scopus in March 2025 we obtained the number of citations.

*Note: The X API was used to have a record of the number of times the institutional account (@INC_UAB) tweeted or retweeted an article. According to this data, the in-house account posted original tweets for 27 of the 201 articles (11 PR and 16 NPR) and 19 of these were not only tweeted but also retweeted (10 PR and 9 NPR). Other 25 articles had from 1 to 8 retweets (RTs) from @INC_UAB, without any original tweets from this account (2 PR and 23 NPR). Finally, 149 articles did not receive any mentions from @INC_UAB (15 PR and 134 NPR)*.

### 2.3 Prospective experiment

The third investigation of this study consisted in the analysis of the prospective outcome of in-house neuroscience publications when undergoing controlled dissemination campaigns. During 2022 and 2023, 25 original articles from members of our center as principal authors were selected and divided into two groups: those for which we would deliver a PR (PR’ group, n=12) and those for which we would not (NPR’ group, n=13). The division was made considering a fork decision tree that importantly took into account 1) whether the article would be of general interest, 2) the preference of the author/s, and 3) the time lapse between the publication of the article and the start of the communication campaign (**Fig. 3**).

**Fig. 3.**
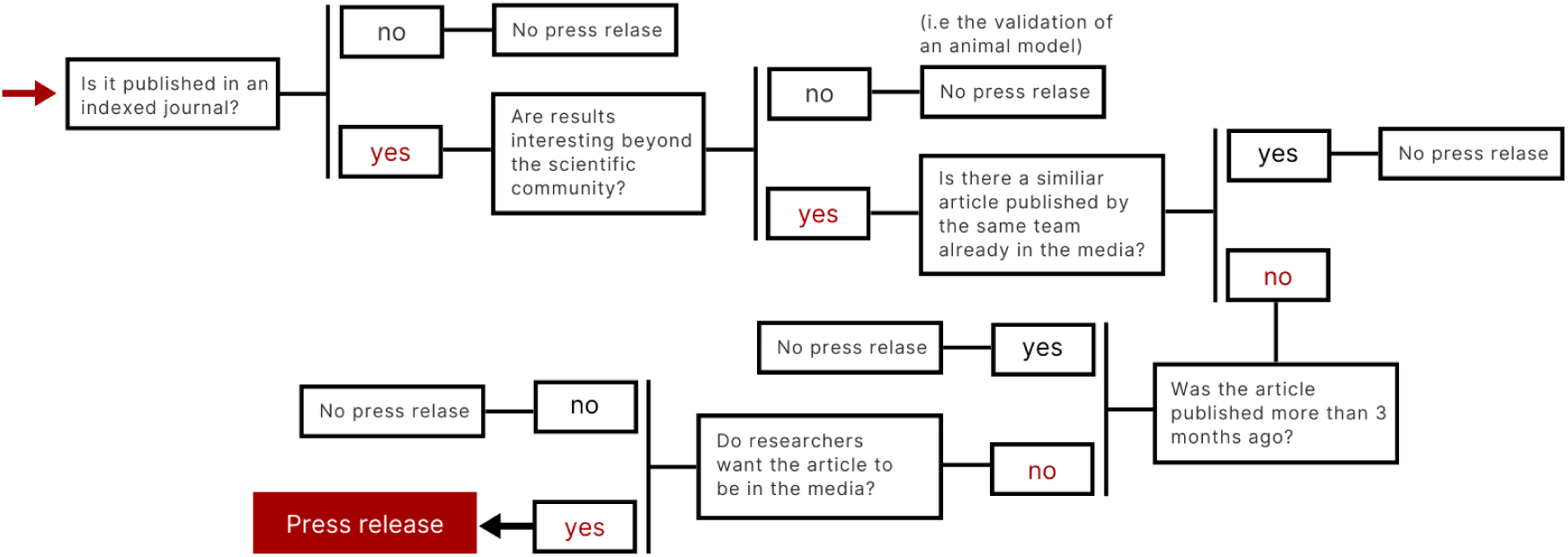
Decision tree on the choice of sending a press release about a scientific article. *Note: the three-month limitation is due to Eurekalert! policy only allowing posting news about publications within the last 90 days*.

*Note: We acknowledge that the distinction between what is considered interesting beyond the academic community and what is not is inherently subjective. In our analysis, we categorized articles as not newsworthy if they focused on laboratory protocols, the validation of animal models, or highly specific biochemical pathways without clinical implications, for example. Although these criteria may introduce bias, the selected articles we studied were compared to themselves, before and after the communications efforts, allowing each publication to serve as its own control. Therefore, any observed differences in attention metrics reflect the impact of the dissemination campaign. That said, further research to better define what truly captures public interest would be highly valuable*.

Within the PR’ and the NPR’ groups, we applied another subdivision: those articles for which we would prepare dissemination materials for publication in SM (sm articles), particularly on X and Instagram (IG) (PR’sm and NPR’sm groups), and those for which we would not (n articles) (PR’n and NPR’n groups). For this subdivision, we also had to consider the preferences of the leading authors. Although the articles in the PR’n and NPR’n groups were intentionally not tweeted, they could be retweeted upon the authors request. These articles remained in the n groups, as such actions were deemed residual (**Table 1**).

**Table 1:** Description of the communication actions carried out for each article. The subscript in the reference indicates the number of days between the article’s publication and the start of the advertising campaign. *Note: The article PR’_68_n corresponds to article n(6’) in graphs 10c, 11c, 12d, and 13*.

| Reference and group | IF | Publication date | Comm. Campaign start | Days of delay | Communications actions by INC-UAB |
| --- | --- | --- | --- | --- | --- |
| PR<sub>0sm</sub> | 25 | 16.06.2022 | 16.06.2022 | 0 | PR + 1 X thread on the same day of the article publication. 7 RTs the day after. 1 RT one week after. 1 tweet 53 days after (summer reminder). |
| PR<sub>1sm</sub> | 3.7 | 08.05.2023 | 09.05.2023 | 1 | PR + 1 X thread and 1 RT the day after the publication. |
| PR<sub>8n</sub> | 6.2 | 30.06.2022 | 8.07.2022 | 8 | PR 8 days after the publication. |
| NPR<sub>0n</sub> | 16.6 | 28.07.2022 | 28.07.2022 | 0 | 1 RT 100 days after the publication. |
| NPR<sub>0n'</sub> | 7.9 | 11.06.2022 | 11.06.2022 | 0 | 1 RT 2 days after the publication. |
| PR<sub>38sm</sub> | 6.1 | 08/02/2022 | 18/03/2022 | 38 | PR + 1 tweet and 1 RT 38 days after the article publication. 2 RTs the day after (day 39). |
| PR<sub>83sm</sub> | 9.3 | 02.02.2022 | 26.04.2022 | 83 | PR + 2 tweets (part of the same thread) and 2 RTs + 1 LinkedIn post 83 days after the publication. 1 RT the day after (day 84). 1 tweet 114 days after (summer reminder). |
| PR<sub>88sm</sub> | 7 | 16/09/2022 | 13/12/2022 | 88 | PR + 1 tweet 88 days after the publication. 1 tweet one week after. |
| PR<sub>41sm</sub> | 5.9 | 24/03/2023 | 04/05/2023 | 41 | PR 41 days after the publication. 1 X thread the day after (day 42). |
| PR<sub>23sm</sub> | 6 | 30/05/2023 | 22/06/2023 | 23 | 2 RTs 1 day after the publication. 1 RT 14 days after the publication. PR + 1 X thread 23 days after the publication. |
| PR<sub>75n</sub> | 6 | 08/01/2022 | 24/03/2022 | 75 | PR 75 after the publication. |
| PR<sub>56n</sub> | 2.6 | 24.03.2022 | 19.05.2022 | 56 | PR 56 after the publication. |
| PR<sub>68n</sub> | 6 | 14/10/2022 | 21/12/2022 | 68 | PR 68 after the publication. |
| PR<sub>64n</sub> | 6.9 | 17/05/2023 | 20/07/2023 | 64 | PR 64 after the publication. |
| NPR<sub>28sm</sub> | 5.3 | 21.09.2022 | 19.10.2022 | 28 | 1 RT one day after the publication. 1 X thread the day 28. 1 RT of the X thread on the day 29 and the day 33. |
| NPR<sub>32sm</sub> | 3.3 | 22.04.2022 | 24.05.2022 | 32 | 1 X post 32 days after the publication (not detected by Altmetric). 1 summer reminder 124 days after. |
| NPR<sub>31sm</sub> | 4.6 | 09.05.2022 | 09.06.2022 | 31 | 1 RT 3 days after the publication. 1 X thread 31 days after the publication. 1 tweet 105 days after (summer reminder). |
| NPR<sub>129sm</sub> | 2.1 | 01.02.2022 | 10.06.2022 | 129 | 1 X thread 129 days after the publication (non detected by Altmetric). 2 tweets 199 days after (summer reminder). |
| NPR<sub>25sm</sub> | 5.6 | 28.11.22 | 23.12.2022 | 25 | 2 tweets (part of the same thread) 25 days after the publication. 1 tweet 28 days after the publication. |
| NPR<sub>75sm</sub> | 2.9 | 22.01.2022 | 07.04.2022 | 75 | 1 X thread 75 days after the publication. 1 tweet 216 days after (summer reminder). |
| NPR<sub>27sm</sub> | 4.1 | 14.10.22 | 10.11.2022 | 27 | 2 tweets (contained in 1 X thread) 27 days after the publication. Another X thread on the day 34. |
| NPR<sub>52n</sub> | 1.3 | 05.03.2022 | 26.04.2022 | 52 | None |
| NPR<sub>90n</sub> | 5.6 | 01.04.2022 | 03.06.2022 | 90 | None |
| NPR<sub>59n</sub> | 4.7 | 11.04.2022 | 09.06.2022 | 59 | 1 RT 11 days after the publication. |
| NPR<sub>45n</sub> | 7.4 | 11.11.22 | 26.12.2022 | 45 | None |

We also made a third subdivision based on the timing of the communication efforts, in relation to the online publication date of each article. According to Fang & Costas (2020), by day 22 after the publication of an article, half of the mentions in the news, and around 60% of the X posts have already happened. Considering this, we set up two groups taking into account whether these 22 days have passed or not (+22 or −22) before the campaign started. On the one hand, PR’_+22_ and NPR’_+22_ (including PR’_+22_sm, NPR’_+22_sm, PR’_+22_n, and NPR’_+22_n) refer to those articles for which the communication actions took place at least 22 days after their publication (from day 22 onwards). On the other hand, PR’_-22_sm, NPR’_-22_sm, PR’_-22_n, and NPR’_-22_n refer to those for which the communication campaign started before day 22 (from day 0 to 22). Throughout the manuscript and tables, the subscript on each article reference indicates the exact number of days since its publication.

To generate a PR, our communication department worked together with the press office of our university (Universitat Autònoma de Barcelona), and counted on the collaboration of the publication authors (**Fig. 4a**). Our PRs consisted of a summary text of the article, following the National Institute of Health (NIH) recommendations (2018a, 2018b), accompanied by one or two images (**Fig. 5**). PRs were delivered to journalists working in Spanish and Catalan media outlets, and an English version was posted on *Eurekalert!* and *AlphaGalileo*, which are platforms to share research news internationally.

**Fig. 4.**
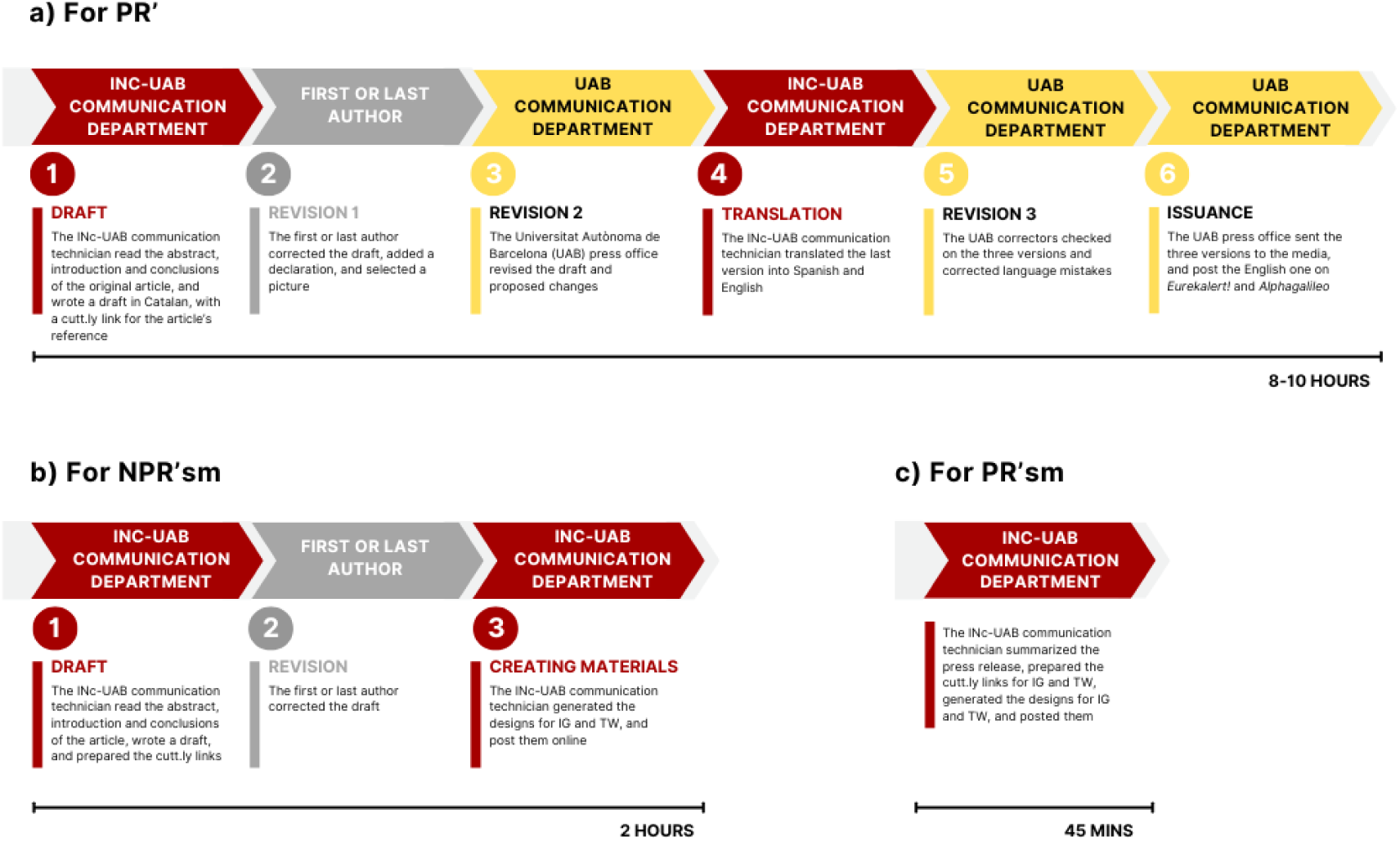
Internal workflows at INc-UAB. **a)** For preparing a PR **b,c)** For generating SM posts about an article.

**Fig. 5.**
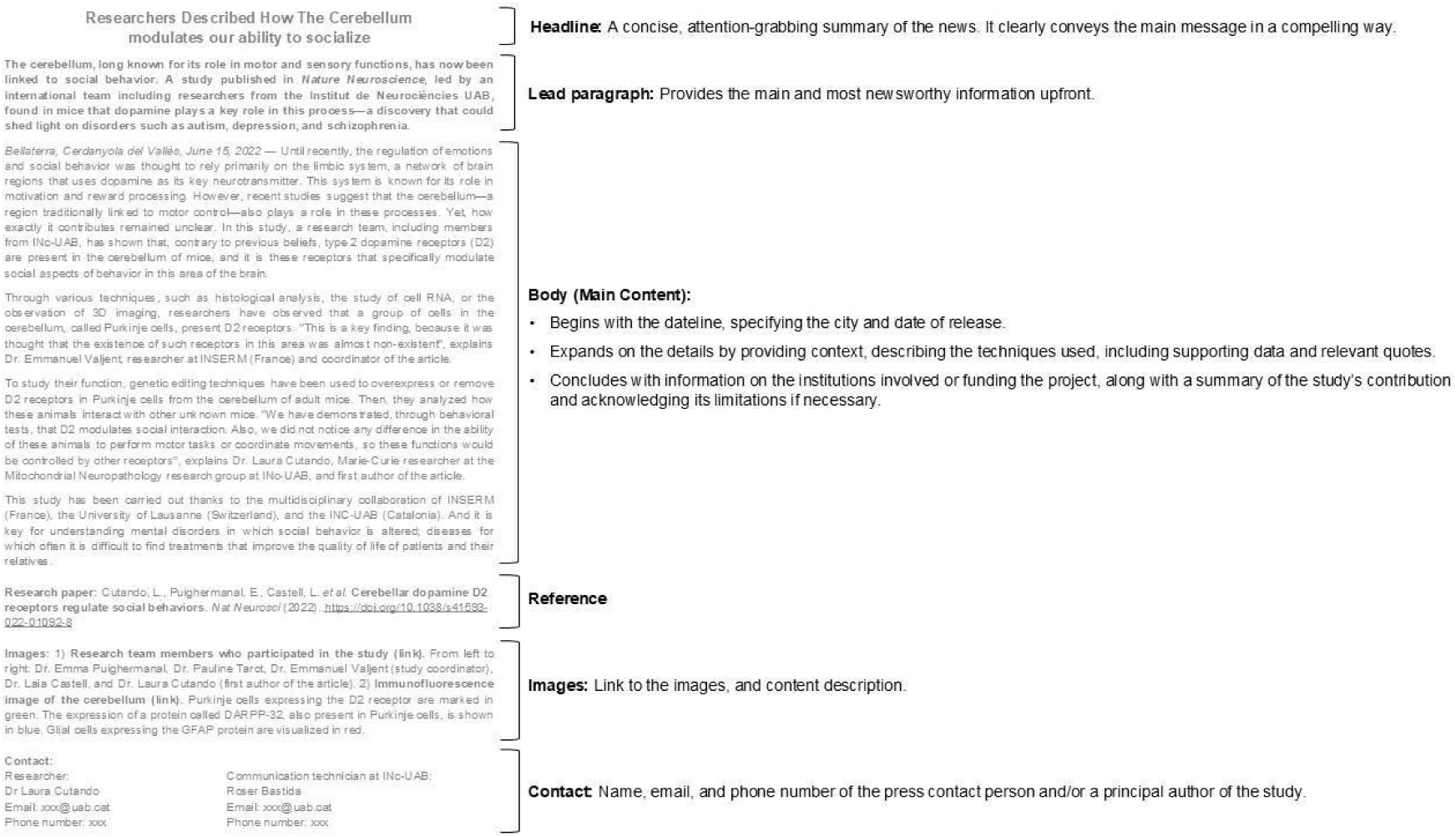
Example of a press release produced by the INc-UAB communication department. Following the standard format used by the UAB, to announce the publication *‘Cerebellar dopamine D2 receptors regulate social behaviors’* (PR’0sm). Once drafted, it was reviewed, at a minimum, by the lead author and the UAB press office, to ensure that the content, structure and language were optimal.

Regarding SM, we used X and not Facebook, as nowadays the usage of Facebook follows a steady decline (Cannarella and Spechler, 2014). Instead, we posted on IG, which at this moment is not tracked by Altmetric but allowed us to engage and communicate with our student community. Materials for X and IG were created with inspiration from the *Three Facts and a Story* method described by Tapper et al. (2021) (**Fig. 4b, c**). The design and posting strategy were as follows: For IG - One carousel post per article with text and images, following the structure: 1–3 images for the introduction, 1–3 to explain the article, and 1–2 to explain the importance of the contribution (see a diagram and an example in **Fig. 6a, b**). On the post text, we included the hashtag #articledeldia (“article of the day”, in Catalan) and a short sentence indicating that the link to the article could be found on the INc-UAB IG profile. The post was shared once as an IG story, tagging the researchers involved.

**Fig. 6.**
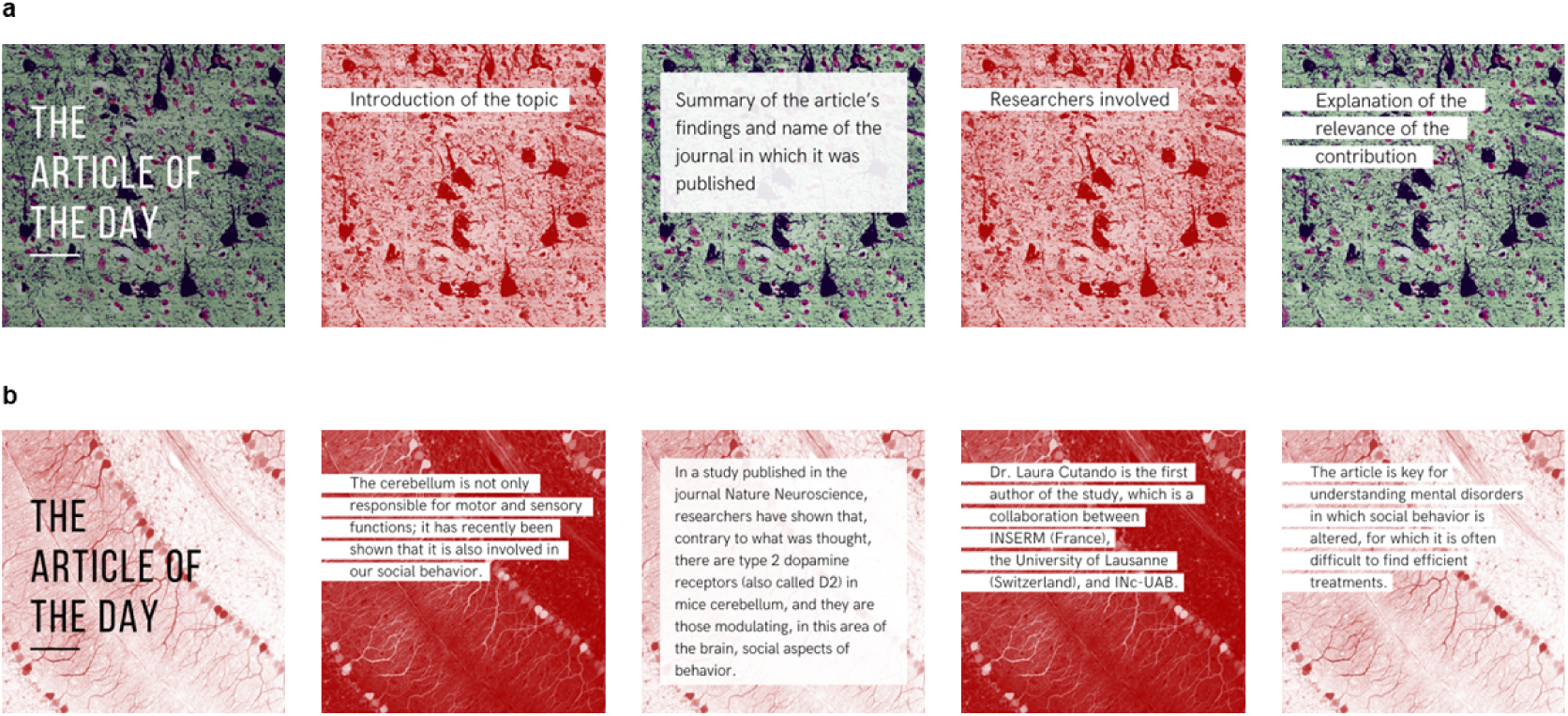
Instagram posts to share the results and contributions of a research article. **a)** Scheme for an IG post **b)** Example of IG carousel for the publication ‘Cerebellar dopamine D2 receptors regulate social behaviors’ (PR’_0_sm): https://www.instagram.com/p/Ce34whGjp1j/?img_index=1

For X - We posted a thread with the same information posted on IG but adapted to X, using the same hashtag #articledeldia (see a scheme and an example in **Fig. 7a, b**). We tagged the journal, authors, labs, and other research centers involved in the study when they had X accounts. Exceptionally, in two cases, we only posted one tweet with all the information instead of a thread. The thread/tweet was shared once and, in some cases, repeated (or a summary of it) three or four days later. Also, for 6 of them (2 in the PR’sm group, and 4 in the NPR’sm group) a reminder was posted a few months later, coinciding with the summer holidays (**Table 1**).

**Fig. 7.**
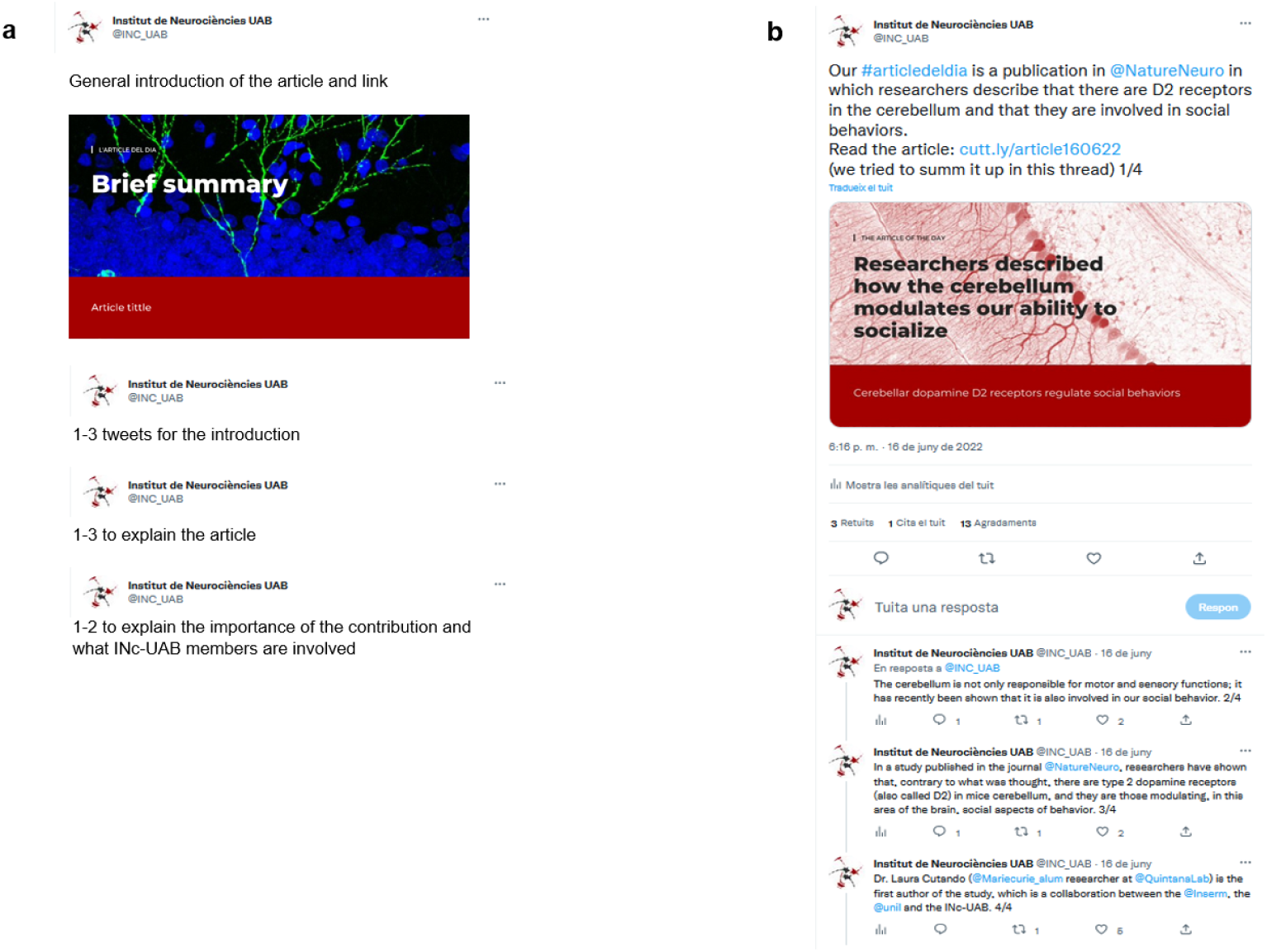
X posts to share the results and contributions of a research article. **a)** Scheme for a X post **b)** Example of a X post for the publication ‘Cerebellar dopamine D2 receptors regulate social behaviors’ (PR’_0_sm): https://x.com/INC_UAB/status/1537469047788425219

One article (PR’_83_sm), was also highlighted on the INc-UAB LinkedIn account, but we stopped using this platform as Altmetric was not providing LinkedIn information at the time when this study was performed, and it was not included in the AAS weighting. As it was said above, IG was not included in the AAS either, but we used it principally as an internal communication channel with our students.

Data was collected six times along the timeline: just before the communication campaign started, and one day, three days, one week, one month, and three months after the communication campaign. We used the AAS explorer, the X searcher, and Google to gather the data. Tweets and retweets were counted as one X post. Data regarding the days before the start of the communication campaign was calculated afterwards, using the Altmetric Explorer and the indicative score table provided by Altmetric (Altmetric, 2023).

The number of abstract and full-text reads was only possible to obtain in those cases in which the journal offered the information. The number of clicks on SM posts or PRs was followed on cutt.ly. Finally, the analysis of the news items, which included checking their language and whether they properly referenced the article, was conducted in August 2025. In some cases, the full text was no longer accessible; however, since the title had been recorded beforehand, we were still able to determine the language.

### Statistical analysis

For the first and second studies, we used Pearson coefficients for correlations and the t-student test for comparing two groups. Partial correlations and ANCOVA tests were also performed to control the effect of particular variables. When homogeneity of variance was not achieved, logarithmic transformations were performed or we used non-parametric tests (Spearman). In study 2, for the matched analysis, a KDTree algorithm (implemented in Python) was used to find the nearest IF in each case, ensuring that each match was unique and avoiding duplicates. Then a paired t-test was performed. For study 3, comparisons between articles with and without PR, and with and without SM campaigns, were conducted using ANOVA and ANCOVA.

Because the number of citations and the number of Mendeley readers increase with the years (Schloegl & Gorraiz, 2011), for correlations involving citations and Mendeley readers (study 1 and 2), these values were normalized, dividing them between the average number of citations or Mendeley readers of the year. This operation allowed the comparison between articles and reviews published in different years.

For this study, all other variables were considered to be equivalent among the years, as their main accumulation usually is very fast within the first few days after the publication (Fang & Costas, 2020).

In study 3, the variable TIME refers to the temporal factor within the statistical analysis. Specifically, it represents the variation of the PR or SM effect on the altmetrics variables over time.

Analyses were performed using Statistical Package for the Social Sciences (IBM SPSS Statistics 29.0.2.0), and graphics were created with GraphPad Prism 8.0.1. Statistical significance was assessed as p<0.05.

## 3. RESULTS

### 3.1 Retrospective study of articles and reviews published by 8 worldwide Neuroscience Research Centers

To have a clear picture of the factors that increase the impact of a publication in the field of neuroscience, in this part of the study, we examined the existing correlations between social attention and citations, using a sample of over 3000 publications of internationally recognized neuroscience centers.

#### 3.1.1 A higher social impact correlate with increased citations

We performed Pearson tests to evaluate the potential correlation between bibliometric indexes (citations) and alternative metrics. Either in the case of original articles or reviews, citations significantly correlated with X posts, Facebook posts, mentions in the news, Mendeley readers, AAS, and AAS percentile (**Table 2**).

**Table 2.** Pearson correlations between citations and X posts, Facebook posts, mentions in the news, Mendeley readers, AAS and AAS percentile, for articles (n=2745) and reviews (n=384), with all variables log-transformed. **Significance at the p<0.01 level.

|  | X Posts | Facebook posts | Mentions in the news | Mendeley readers | AAS | AAS percentile |
| --- | --- | --- | --- | --- | --- | --- |
| Articles | 0.49** | 0.23** | 0.42** | 0.78** | 0.50** | 0.23** |
| Reviews | 0.44** | 0.23** | 0.45** | 0.84** | 0.50** | 0.23** |

As journal quality may also influence the number of citations or altmetrics, we examined the correlation between IF, citations, and AAS, and found significant associations. IF correlated with the number of citations, with coefficients of 0.55** for articles and 0.51** for reviews, and with AAS, with coefficients of 0.49** for articles and 0.43** for reviews (p < 0.01). Consequently, all previous correlations were recalculated adjusting for IF, and we found that alternative metrics could still predict scholarly impact (**Table 3**).

**Table 3.** Pearson partial correlations between alternative metrics and number of citations, after controlling for IF, with all variables log-transformed. Significance at the *p<0.05 level or **p<0.01 level.

|  | X Posts | Facebook posts | Mentions in the news | Mendeley readers | AAS | AAS percentile |
| --- | --- | --- | --- | --- | --- | --- |
| Articles | 0.31** | 0.15** | 0.25** | 0.71** | 0.32** | 0.12** |
| Reviews | 0.28** | 0.15* | 0.41** | 0.77** | 0.37** | 0.18** |

#### 3.1.2 X is the social network most widely used to mention neuroscience articles

According to the Altmetric API, more than 90% of the articles published by these centers were mentioned at least once on X. However, mentions on Facebook only reached 28.2%, and only 24% on the news (**Table 4**). Furthermore, analyzing SM per year, we found that in 2020, 92.1% of the publications appeared on X at least once, while only 17.9% did on Facebook, with a clear decay of the latter from 2018 onwards (**Fig. 8a**).

**Fig. 8.**
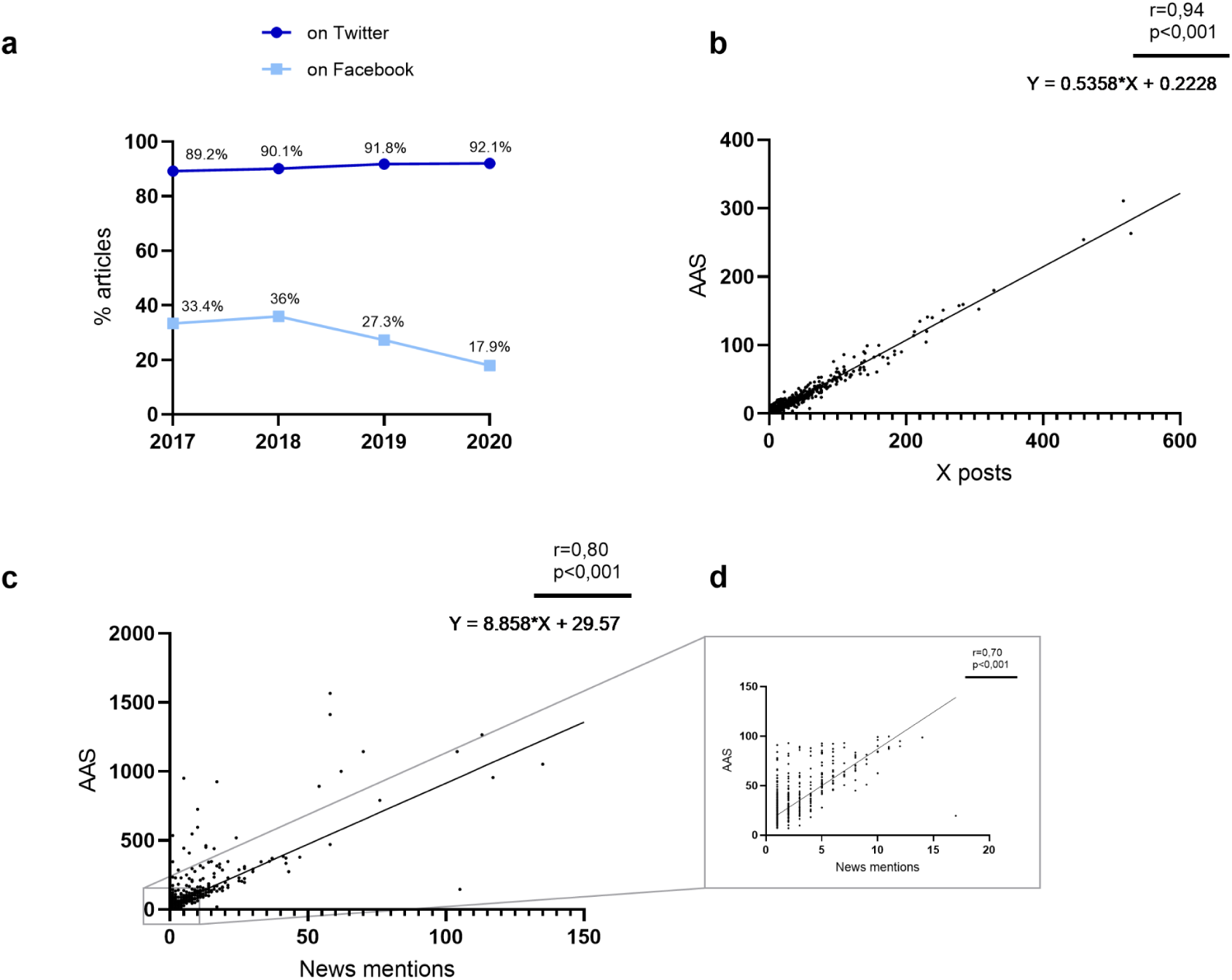
X is the social platform most frequently utilized to cite neuroscience articles, but press mentions have the greatest impact on increasing AAS. According to Altmetric data, **a)** Evolution of the number of publications that appeared at least once on X and Facebook per year **b)** Spearman correlation between X posts and AAS for publications not mentioned in the news **c)** Spearman correlation between news mentions and AAS for articles mentioned at least once in the news **d)** Spearman correlation between news mentions and AAS for articles mentioned in the news with an AAS below 100. *Note: While the correlation analysis in the tables 2, 3 and 4 was performed using log-transformed data, these graphs display untransformed data to facilitate interpretation of absolute changes*.

**Table 4.** Percentage of publications mentioned on SM or in the news, categorized by the number of mentions.

|  |  | Number of times the publication is mentioned |  |  |  |  |  |  |
| --- | --- | --- | --- | --- | --- | --- | --- | --- |
|  |  | 0 | 1 | 2 | 3 | 4 | 5 | >5 |
| Articles | X | 9.7% | 10.9% | 7.7% | 6.7% | 5.6% | 5.2% | 54.2% |
|  | Facebook | 71.9% | 16.8% | 6% | 2.7% | 1% | 0.5% | 1.1% |
|  | News | 76% | 11.9% | 2.9% | 1.2% | 1.1% | 1% | 5.9% |
| Reviews | X | 6 | 6.8% | 10.9% | 4.7% | 4.2% | 5.5% | 62% |
|  | Facebook | 68.2% | 15.1% | 7.6% | 3.4% | 3.1% | 0.3% | 2.3% |
|  | News | 75% | 13% | 3.6% | 1.6% | 0.8% | 0.8% | 5.2% |

On average, X is where more mentions appeared. Interestingly, though, while AAS increased by around

0.5 points for each tweet or retweet (**Fig. 8b**), it increased by more than 8 points for each mention in the news (**Fig. 8c, d**). These results, which are in line with what is predicted by Altmetric (Altmetric, 2023), suggest that Altmetric considers a mention in the press may result in a much greater impact than in SM. Moreover, not only the type of source and number of mentions matter: this relation is not linear. Altmetric also takes into account factors such as the popularity of the media outlet or author, as well as their posting patterns, which influence the score assigned (Altmetric, 2024).

### 3.2 Retrospective study of in-house articles

In our previous analysis described above, we did not have information on the nature of the communication campaign for each case. To fill this gap, we conducted a second study focused on in-house articles, analyzing 201 publications with known campaign details. The aim was to examine whether active communication efforts are associated with enhanced altmetric performance, as reflected in the number of X posts, news mentions, and the resulting AAS.

#### 3.2.1 Retrospective analysis indicates that articles with a press release have stronger altmetric performance

Since previous studies suggest that PRs greatly improve media attention (Cho & Yoon, 2019; de Semir et al., 1998; Fuoco et al., 2023; Stryker, 2002), we compared those in-house articles for which we delivered a PR (PR group) to those for which we did not (NPR group). We found that the PR group (n=28) not only received 39.5 times more mentions in the news (**Fig. 9a**) but experienced a ripple effect reflected in 3.7 times more X mentions (**Fig. 9b**) and 1.6 more Mendeley readers (**Fig. 9c**). PR articles had an AAS average of 76.3 points, whereas the NPR (n=173), of 4.58, with a difference higher than 70 points. This represents a 16.6-fold difference (**Fig. 9d**), indicating that delivering a PR may be a critical action to increase the general online social attention of an article. Notably, this effect was observed across all IF ranges (**Fig. 9e**).

**Fig. 9.**
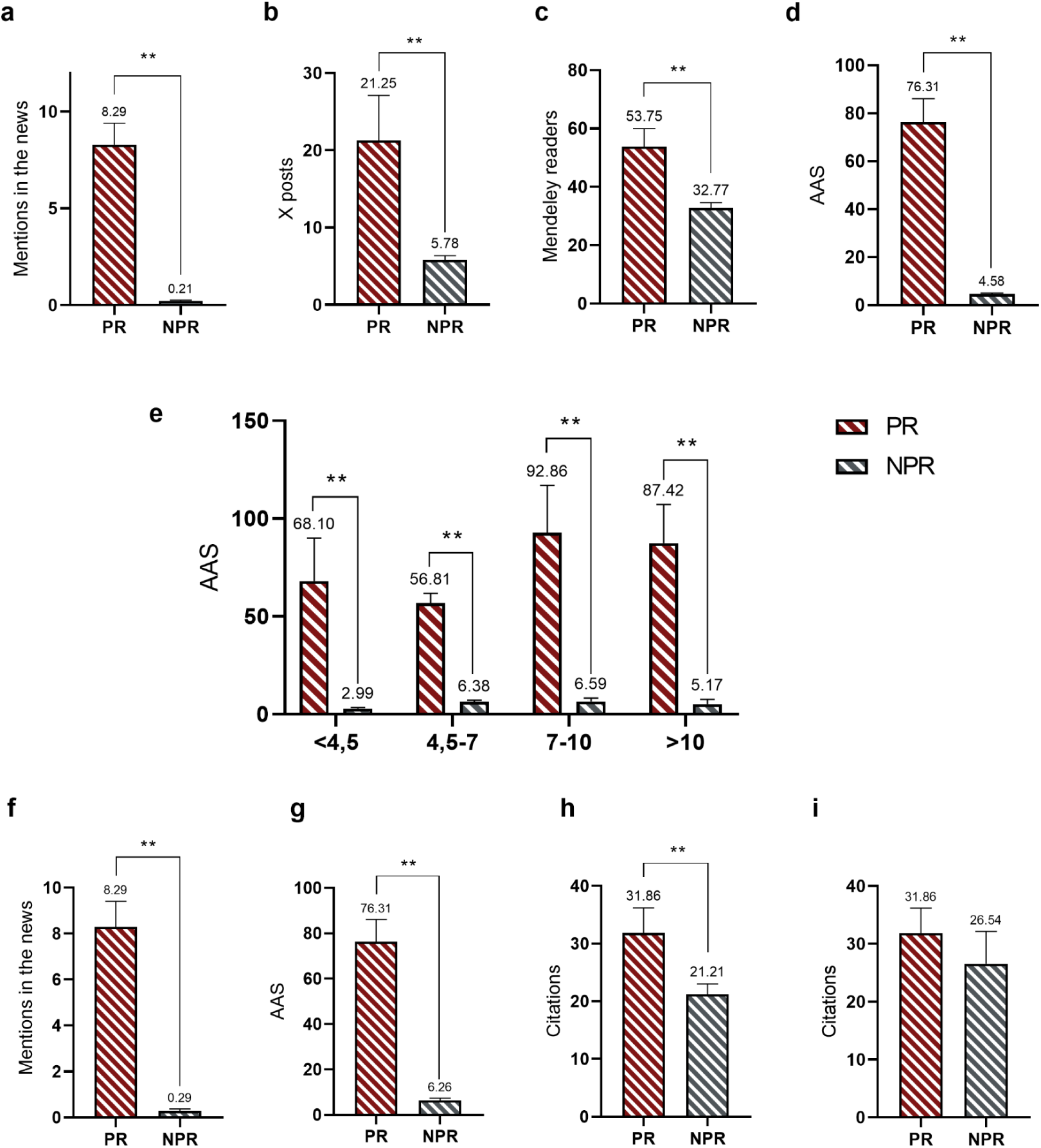
PR articles have higher altmetrics and citations, although the citation advantage disappears after controlling for journal IF. Comparison between PR and NPR groups regarding **a)** The number of mentions in the news according to Altmetric [t(199)=-25.043, p<0.01] **b)** The number of X posts according to Altmetric [t(199)=-6.263, p<0.01] **c)** The Mendeley readers according to Altmetric [t(199)=-4.457, p<0.01] **d)** The AAS [t(199)=-15.541, p<0.01] **e)** The AAS, per IF: <4.5 [t(97)=-9.274, p<0.01], 4.5-7 [t(69)=-6.359, p<0.01], 7-10 [t(16)=-5.948, p<0.01], >10 [t(11)=-5.506, p<0.01] **f)** The number of mentions in the news in a matched analysis by IF [t(27)=-14.364, p<0.01] **g)** The AAS in a matched analysis by IF [t(27)=-11.171, p<0.01] **h)** The number of citations according to Scopus [t(199)=-2.783, p<0.01] **i)** The number of citations in a matched analysis by IF [t(27)=-1.11, p=0.276]. Means and SEM are represented. *Note: While the graphical representation utilizes untransformed data for clarity, the underlying statistical analyses (t-tests) and their interpretations are based on the logarithmically transformed datasets*.

Given that the first study of the present article found a correlation between journal IF and higher alternative metrics, we controlled for the effect of IF when assessing the influence of the PR. An ANCOVA test showed that the PR effect remained highly significant for the number of X posts [F(1,198)=20.720, p<0.01] and Mendeley readers [F(1,198)=14.531, p<0.01], while it was not possible to calculate for the number of mentions in the news due to the NPR group having not enough variance. Since this was the case, a matched analysis comparing the mentions in the news between PR and NPR articles with similar IFs was performed, showing that significant differences between the two groups persisted [t(27)=-14.364, p<0.01] (**Fig. 9f**). On the other hand, the effect of the PR after controlling for the IF was also significant for the AAS [F(1,198)=178.740, p<0.01], and this was also confirmed by a matched analysis [t(27)=-11.171, p<0.01] (**Fig. 9g**).

Finally, regarding traditional bibliometric indexes, we found that the PR group had significantly higher citations (**Fig. 9h**), although this effect was no longer maintained after adjusting for IF [F(1,198)=2.339, p=0.128] and was also absent in the matched analysis [t(27)=-1.11, p=0.276] (**Fig. 9i**). This suggests that the quality and reputation of the journal are the primary factors influencing academic citations, regardless of social attention.

#### 3.2.2 X posts moderately influenced media coverage but to a lesser extent

Since PR articles generated a ripple effect in X attention, we aimed to test whether this phenomenon could occur in the opposite direction. To do so, we analyzed articles without a PR (NPR) and, after controlling for IF, we found that while PR exponentially increases X mentions, the reverse effect (X posts leading to more news mentions) was smaller but still significant [F(26,145)=1.823, p<0.05]. In addition, X mentions also had a significant impact on Mendeley readers [F(26,145)=2.538, p<0.01], whereas their influence on citations was very weak [F(26,145)=1.547, p=0.057].

### 3.3 Prospective experiment

Given that the in-house retrospective analysis suggests that a PR could be a key factor in boosting the altmetrics of a publication, in this part of the study we test its real impact by comparing the metrics before and after the dissemination campaign.

#### 3.3.1 Mentions in the news only happened for articles with a press release

After waiting at least 22 days after the publication of the article, the communication campaign (consisting of SM posting or issuing a PR) started. Articles for which a PR was distributed (PR’_+22_, n=9) began receiving mentions in the news just hours after the issuance, whereas articles without a PR (NPR’_+22_, n=11) received none. According to Altmetric, and after excluding false positives (records from *Alzforum, Eurekalert!* and *AlphaGalileo,* which are not news items), within the PR’_+22_ group, those that were highlighted on SM (PR’_+22_sm, n=5) showed the highest number of mentions, reaching an average of 9.2 per article, while those that were not posted on SM (PR’_+22_n, n=4) averaged 4 mentions, one week after publication (**Fig. 10a–c**; **Table 5**).

**Fig. 10.**
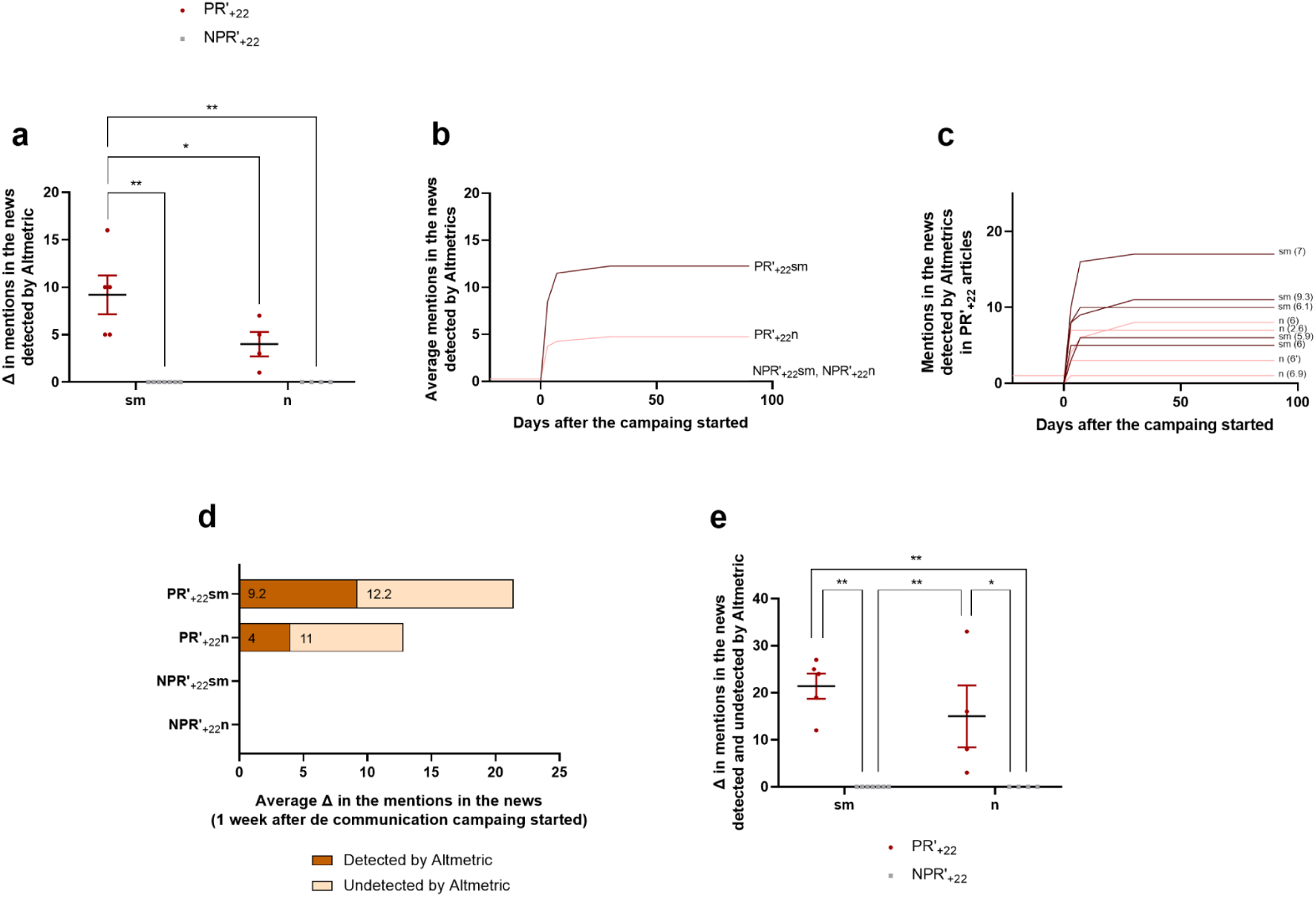
A press release directly affects the number of mentions in the news. According to Altmetric data, after excluding false positives, **a)** Differences among groups regarding their increase in the number of mentions in the news one week after the communication campaign/monitoring started **b)** Average increase in the number of mentions in the news per group, starting at day −22 before the communication campaign/monitoring started, and until 90 days after **c)** Increase in the number of mentions in the news in PR’_+22_ articles (in brackets, the article IF) **d)** Average increase in the number of mentions in the news per group, one week after the communication campaign/monitoring started, detected and undetected by Altmetric **e)** Differences among groups regarding their increase in the number of mentions in the news detected and undetected by Altmetric, one week after the communication campaign/monitoring started.

**Table 5.**
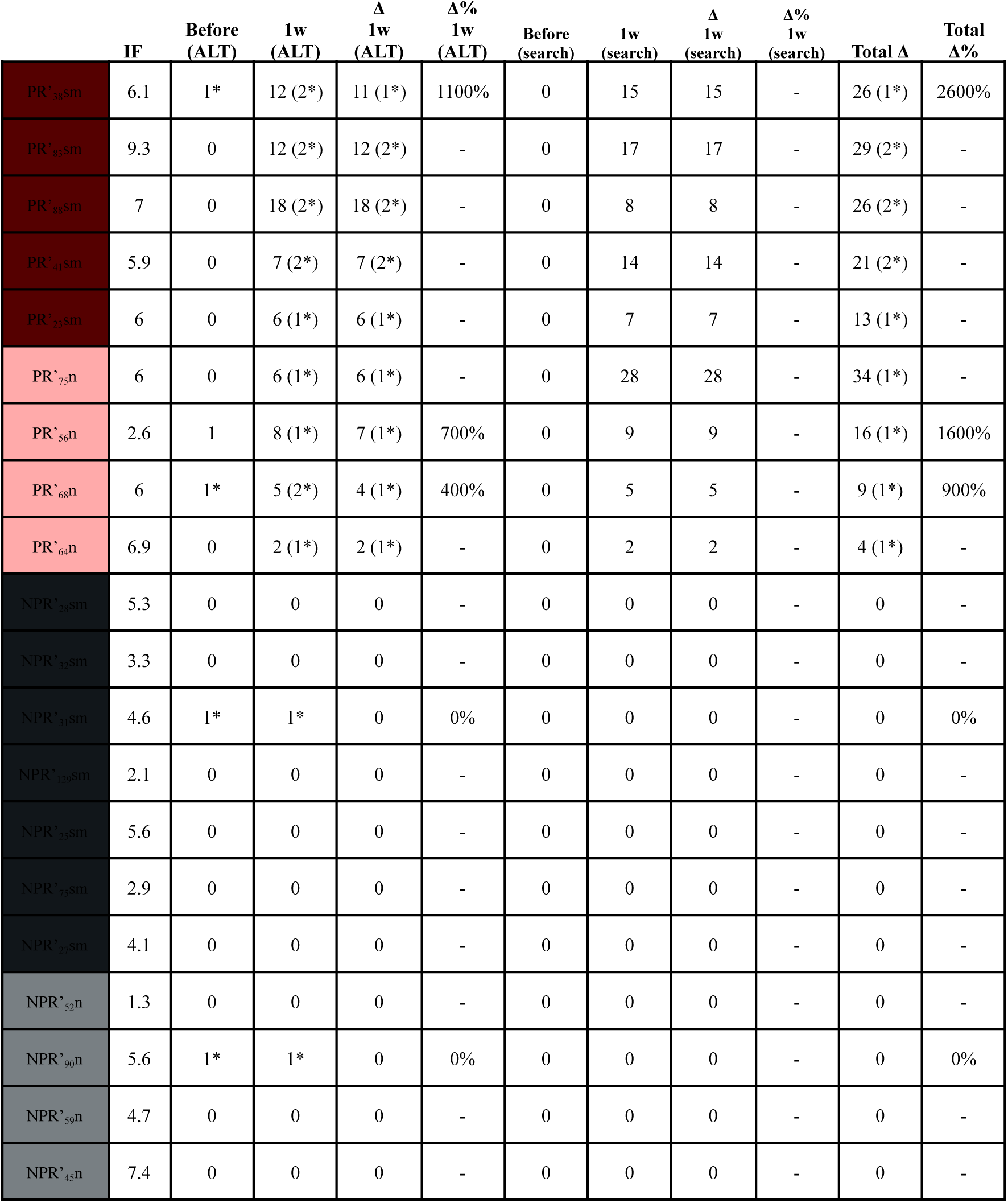
Mentions in the news before and one week after sending a PR (for PR’_+22_sm and PR’_+22_n articles), posting the article in the institutional SM accounts (for PR’_+22_sm and NPR’_+22_sm articles), or starting the monitoring process for the control group (NPR’_+22_n). Data includes mentions detected by Altmetric and those undetected, gathered via manual searches. The subscript in the reference for each article indicates the number of days between the article’s publication and the start of the advertising campaign. *Notes: The article PR’68n corresponds to article n(6’) in graphs 10c, 11c, 12d, and 13. The mentions marked with an asterisk (*) are not pieces of news, although Altmetric would count them as such, but rather a citation in Alzforum or the PR itself posted on the platforms Eurekalert! or AlphaGalileo by the communication office of the University. These false positives were excluded from the statistical analyses and figures*.

However, these numbers were actually underestimated, as Altmetric does not capture a considerable number of news sources. Using Google search, we found that 105 news mentions were overlooked, representing 63% of the total (**Table 5 and 6**). Among the undetected pieces, 10% contained an article reference or a link, 70% did not, and 21% could not be verified as the piece was no longer available. On the other hand, among the detected pieces, 59% contained an article reference or a link, 11% did not, and 30% could not be checked (**Table 7**). Interestingly, when counting detected and undetected news items, 55% of the pieces that could be verified lacked a reference.

**Table 6.**
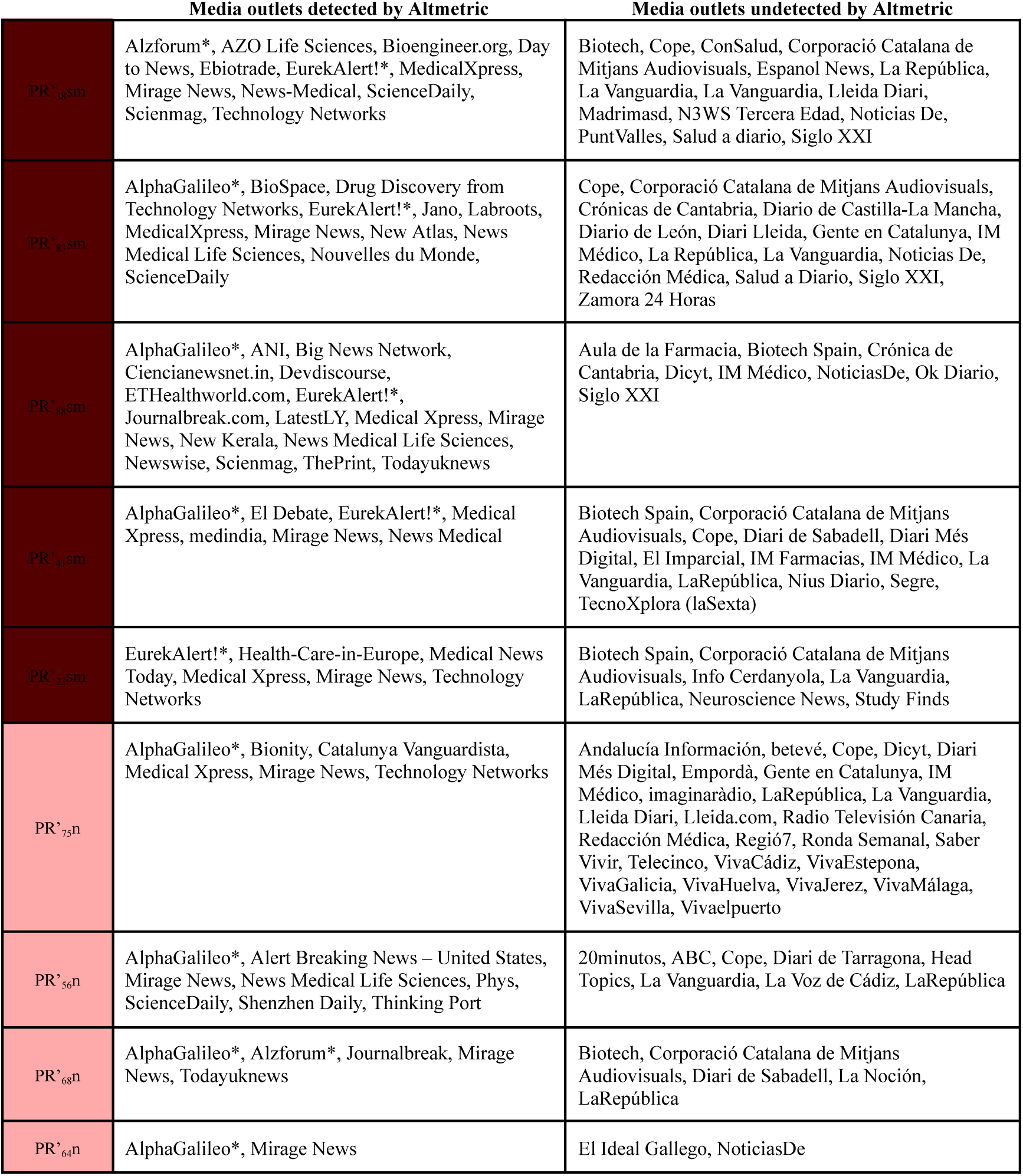
News outlets mentioning each PR article from the prospective study, classified by if they were detected through Altmetric or identified independently through manual searches. *Notes: One media outlet may have more than one news item about one single article (not a translation, but a different piece). The outlets marked with an asterisk are not media platforms, although Altmetric would count them as such. These false positives were excluded from the statistical analyses and figures*.

**Table 7.** Distribution of the pieces of news detected and undetected by Altmetric, according to language and the presence of a reference or link to the publication within the text.

|  | English | Spanish | Catalan | Other |
| --- | --- | --- | --- | --- |
| Detected – referencing | 40 | 4 | 0 | 1 |
| Detected – not referencing | 6 | 2 | 0 | 0 |
| Detected – unknown | 18 | 4 | 0 | 1 |
| Undetected – referencing | 2 | 8 | 0 | 0 |
| Undetected – not referencing | 0 | 47 | 26 | 0 |
| Undetected – unknown | 0 | 17 | 5 | 0 |

After adding the undetected mentions to those already recorded by Altmetric, we observed that within one week, PR’_+22_sm articles received an average of 21.4 mentions per article; PR’_+22_n, of 15, and NPR’ groups did not receive any (**Fig. 10d, e**), (**Table 5**).

Regarding the factors that influenced the differences among groups, after adjusting for IF, the analysis of covariance showed that the interaction between TIME (the moment at which mentions were counted) and PR [F(1,15)=149.204, p<0.01] was statistically significant, while all the other interactions were not [TIMExSM, PRxSM, TIMExPRxSM]. This suggests that the PR is the primary factor that drives the increase in news mentions. In line with this, the decomposition of interactions showed that, prior to the intervention, group differences in the number of news mentions were not statistically significant, whereas PR had a significant effect afterward [F(1,15)=159.832, p<0.01]. SM had no such effect, indicating that the PR is the key factor affecting news mentions.

#### 3.3.2 Press releases increase the number of tweets about an article

When examining the impact of a PR on SM, notable effects were observed. One week after the PR, PR’_+22_sm articles presented a significant increase in X posts (tweets and retweets), with a fourfold rise, averaging 19 posts detected by Altmetric (**Fig. 11a-c**). In contrast, articles with similar SM campaigns but without a PR (NPR’_+22_sm) only experienced a modest increase, with a 1.9-fold rise, averaging 8.9 posts (**Fig. 11a, b, d**), (**Table 8**). On the other hand, when SM campaigns were not applied, articles disseminated with a PR (PR’_+22_n) still showed an increase in the number of X posts, although smaller (1.8-fold, reaching an average increase in 2.5 posts) (**Fig. 11a-c**), while the increase was null (0 posts) for those with neither a PR nor a SM campaign (NPR’_+22_n) (**Fig. 11a, b, d**) (**Table 8**).

**Fig. 11.**
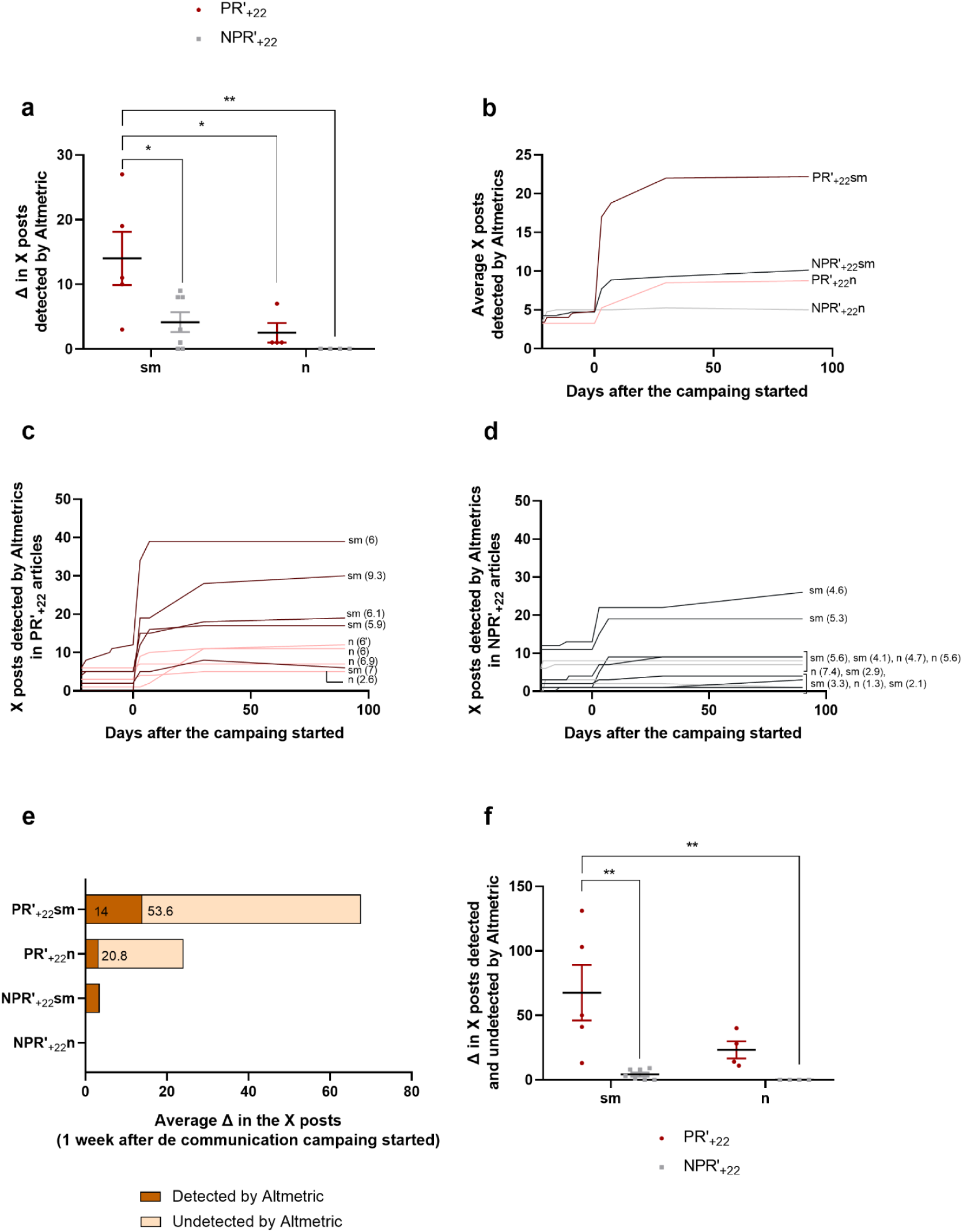
A press release significantly affects the number of mentions on X. According to Altmetric data, **a)** Differences among groups regarding their increase in the number of X posts one week after the communication campaign/monitoring started **b)** Average increase in the number of X posts per group, starting at day −22 before the communication campaign/monitoring started, and until 90 days after **c)** Increase in the number of X posts in PR’_+22_ articles (in brackets, the article IF) **d)** Increase in the number of X posts in NPR’_+22_ articles (in brackets, the article IF) **e)** Average increase in the number of X posts per group detected and undetected by Altmetric, one week after the communication campaign/monitoring started **f)** Differences among groups regarding their increase in the number of X posts detected and undetected by Altmetric, one week after the communication campaign/monitoring started.

**Table 8.**
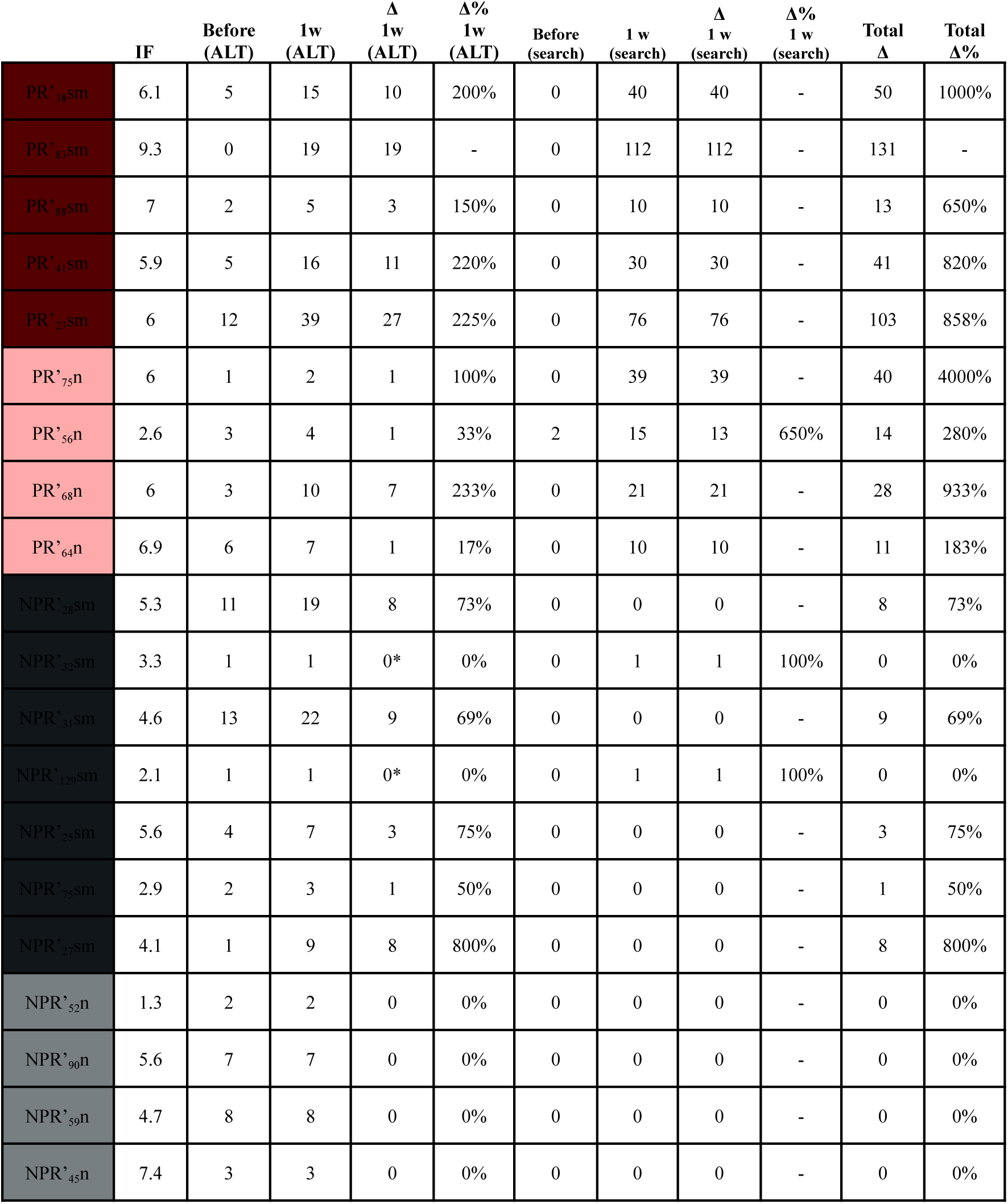
X posts were analyzed before and one week after the PR and SM campaigns, or during the corresponding monitoring period. Data includes those posts detected by Altmetric and those undetected, gathered via manual searches. The subscript in the reference for each article indicates the number of days between the article’s publication and the start of the advertising campaign. *Notes: The article PR’68n corresponds to article n(6’) in graphs 10c, 11c, 12d, and 13. Altmetric did not detect two of our tweets, although they were correctly referenced (cases are marked with an asterisk and are counted in the ‘search’ column)*.

Interestingly, in all groups the number of posts remained stable throughout the whole analyzed timeline. From one week following the beginning of the campaign onward, no significant fluctuations in the number of posts were observed over a 3-month period (**Fig. 11b, c, d**).

As with news mentions, it is important to highlight that Altmetric did not account for a large number of tweets. Specifically, those that did not explicitly cite the publication were excluded from the score, meaning that posts referring to news coverage of the article, rather than the article itself, are overlooked. When using the X search function, we discovered that undetected X posts made up 82% of the total X posts for PR’_+22_ articles but were absent for articles without a PR (**Fig. 11e, f**), (**Table 8**).

When taking into account the posts detected and the undetected, and adjusting for IF, we saw that the interaction TIMExPR was statistically significant [F(1,15)=19.287, p<0.01], while all the other interactions were non-significant [TIMExSM, PRxSM, TIMExPRxSM]. This suggests that the presence of a PR is the main factor underlying the increase in X posts. Moreover, the decomposition of these interactions indicated that, before the intervention, the differences among groups were statistically non-significant, whereas after the interventions, the PR factor was statistically significant [F(1,15)= 16.848, p<0.01]. The number of tweets significantly increased after the PR but not after the SM intervention, indicating that the PR is the key variable responsible for the increase in posts.

Finally, the impact of the number of times each article was tweeted or retweeted by the institutional account cannot be analyzed due to sample limitations, although differences did not seem to be relevant.

#### 3.3.3 Press releases induce an increase of the AAS of an article by almost 70 points

Since the AAS includes both news mentions and X posts, PRs were expected to increase the AAS as well. However, the exact extent could not be predicted because the Altmetric algorithm, which also accounts for source influence and posting patterns, is not fully disclosed (Altmetric, 2023, 2024).

Indeed, while no statistically significant differences were observed between the groups prior to the communication campaign, articles with a PR had a significantly higher AAS post-campaign, even after controlling for IF [F(1,15)=53.676, p<0.01]. As shown on **Fig. 12a-e**, the AAS of PR’_+22_ articles dramatically increased within the first week (68.3 points on average, a 1367%), whereas for the NPR’_+22_ group the AAS only increased by an average of 0.6 points (a 19%).

**Fig. 12.**
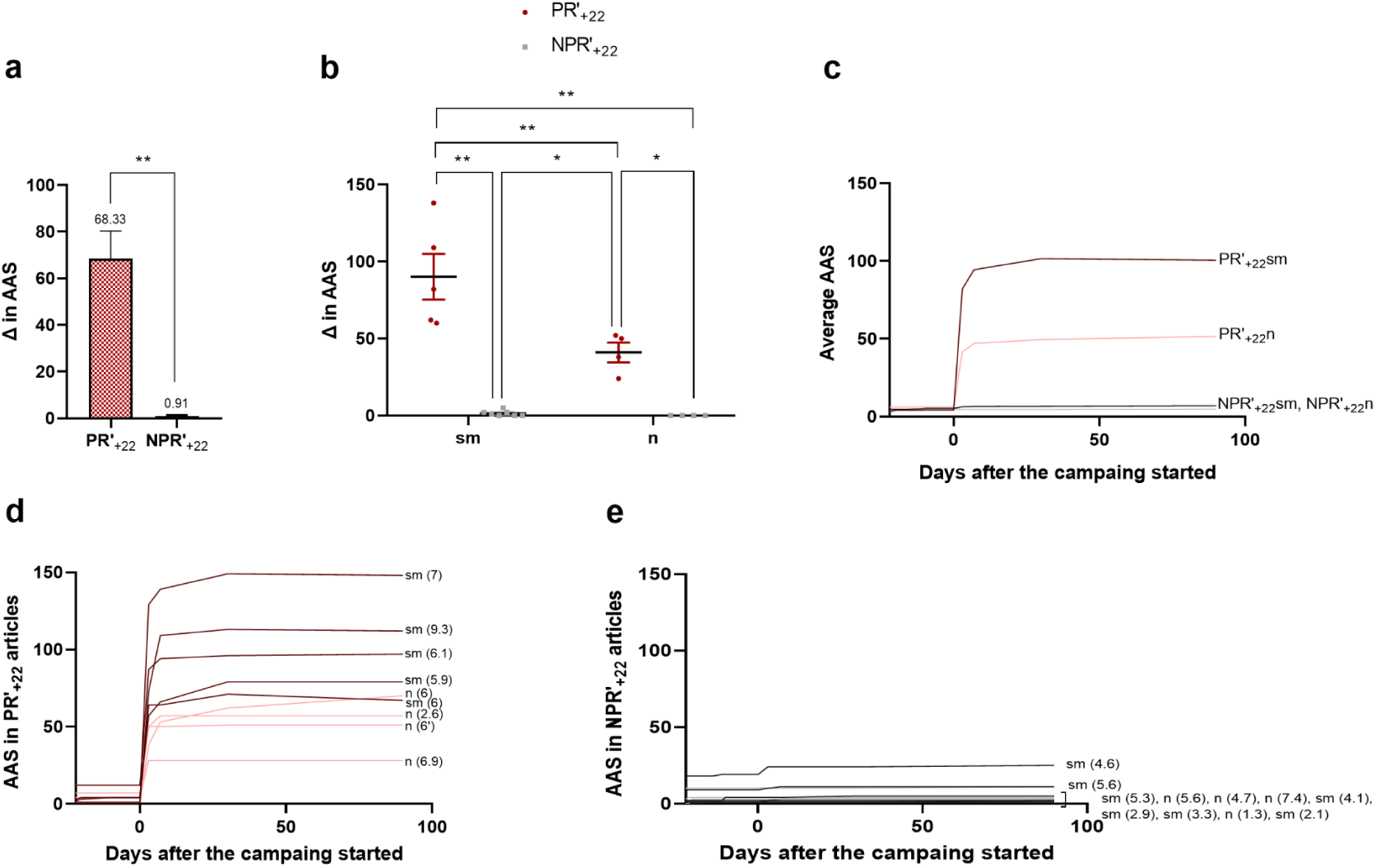
Press release is a critical element of the dissemination campaign to increase online attention. **a)** AAS increase comparison between PR’_+22_ and NPR’_+22_ articles, one week after the campaign/monitoring started [t(18)=**-**14.225, p<0.01]. Means and SEM are represented **b)** Differences regarding the AAS increase considering the presence of a PR or/and SM, one week after the communication campaign/monitoring started (**Significance at the p<0.01 level). *Note: While the a and b graphical representation utilizes untransformed data for clarity, the underlying a and b statistical analyses (t-test and ANCOVA) and their interpretations are based on the logarithmically transformed datasets, ensuring the rigor of our findings.* **c)** Average AAS increase per group, starting at day −22 before the communication campaign/monitoring started, until 90 days after **d)** AAS increase in PR’_+22_ articles (in brackets, the article IF) **e)** AAS increase in NPR’_+22_ articles (in brackets, the article IF).

Regarding the factors influencing these changes, the analysis of covariance showed that the interaction between TIME and PR [F(1,15)=43.668, p<0.01] was statistically significant, while interactions involving SM were not. This indicates that PR is the main variable driving the AAS increase.

Interestingly, the AAS was stable within the rest of the analyzed timeline; from one week after the delivery of the PR or after posting the X and IG materials onwards, showing no significant variations within a 3-month window (**Fig. 12c, d, e**), (**Table 9**).

**Table 9.**
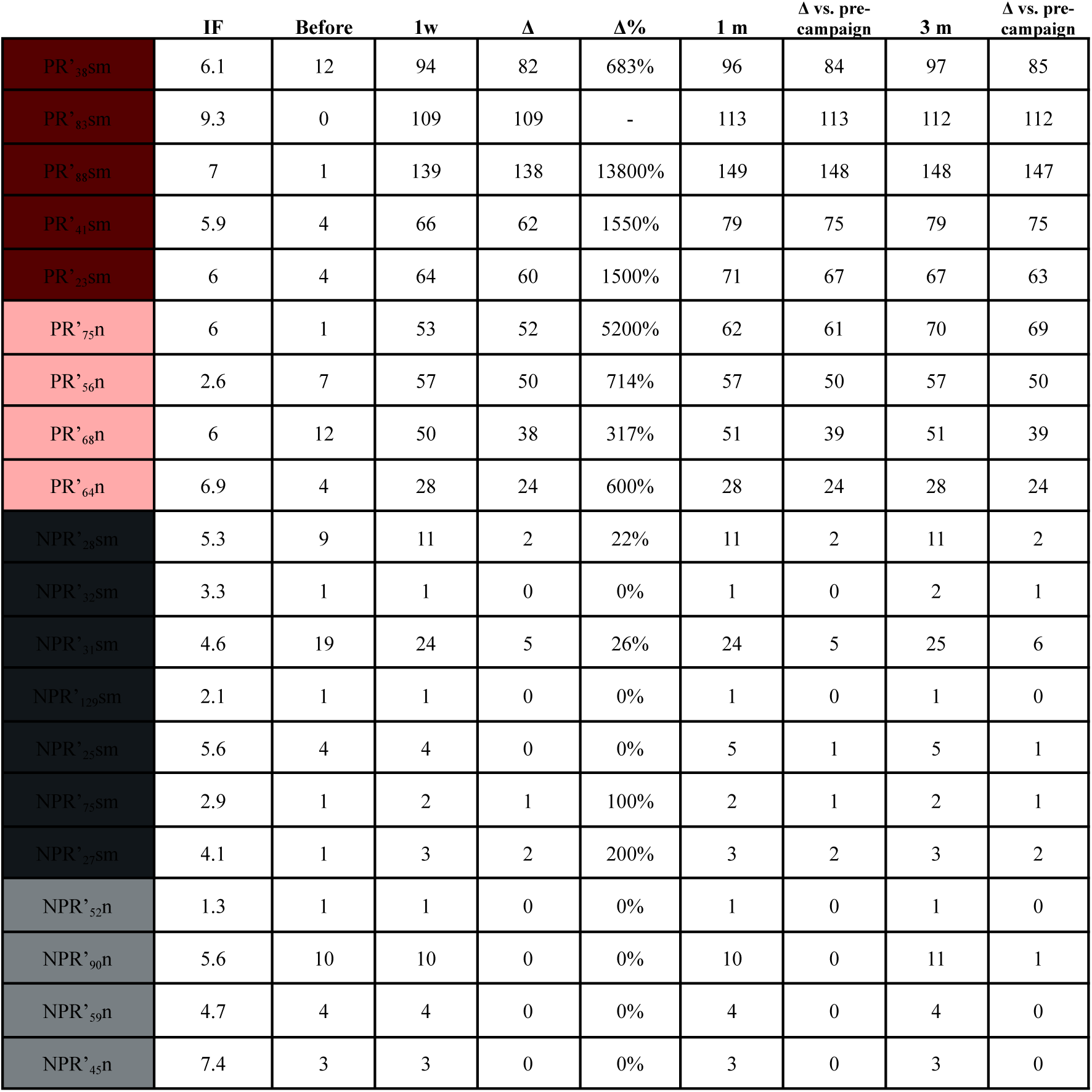
AAS increases before and one week, 1 month, and 3 months after the PR and SM campaigns, or the corresponding monitoring period. The subscript in the reference for each article indicates the number of days between the article’s publication and the start of the advertising campaign. *Note: The article PR’68n corresponds to article n(6’) in graphs 10c, 11c, 12d, and 13*.

It is important to note that the AAS of an article does not always increase over time, it can sometimes decrease (e.g. the AAS of the PR’_23_sm article is higher after 1 month than after 3 months, as shown on **Table 9**). This may occur due to Altmetric readjustments or because some posts or mentions contributing to the initial score are deleted.

As it may also be worthwhile, we examined articles without a 22-day delay between the publication and the communication campaign or monitoring (PR’_-22_ and NPR’_-22_). Among these, two publications with very high IF had distinctly different campaigns, driven by the researchers preferences rather than any assessment of general interest. For one of them (IF=25), a PR was delivered and dissemination materials were posted on SM. For the other (IF=16.6), no communication actions were carried out. In the first week, the first one was mentioned 28 times in the news (17 detected and 11 undetected by Altmetric), 315 times on X (178 detected and 137 undetected), and reached an AAS of 219. However, the other only increased its AAS by 1 point in the first 3 months after its publication, with only two tweets mentioning it after this time (**Table 10)**. This observation, although anecdotal, also points to the repercussions of delivering a PR to significantly increase the AAS regardless of the reputation of the journal where the article is published.

**Table 10.** Mentions in the news, on X, and AAS evolution for articles that did not wait 22 days before starting a communication campaign or monitoring. Changes in the number of mentions in the news or X are reported for one week after the communication campaign/monitoring, while AAS are reported for one week, one month, and three months after the intervention. The subscript in the reference for each article indicates the number of days between the article’s publication and the start of the advertising campaign.

|  | IF | News<br>before | News<br>1w<br>(ALT) | News<br>1w<br>(search) | X<br>before | X<br>1w<br>(ALT) | X<br>1w<br>(search) | AAS<br>Before | AAS<br>1w | AAS<br>1m | AAS<br>3m |
| --- | --- | --- | --- | --- | --- | --- | --- | --- | --- | --- | --- |
| PR'.<sub>0sm</sub> | 25 | 0 | 17 | 11 | 0 | 178 | 137 | 0 | 219 | 225 | 225 |
| PR'.<sub>1sm</sub> | 3.7 | 0 | 8 | 13 | 9 | 80 | 34 | 5 | 100 | 101 | 96 |
| PR'.<sub>8n</sub> | 6.2 | 0 | 17 | 22 | 11 | 15 | 31 | 7 | 133 | 133 | 132 |
| NPR'.<sub>0n</sub> | 16.6 | 0 | 0 | 0 | 0 | 1 | 0 | 0 | 1 | 1 | 1 |
| NPR'.<sub>0n'</sub> | 7.9 | 0 | 0 | 0 | 0 | 20 | 0 | 0 | 12 | 13 | 14 |

#### 3.3.4 After a 22-day delay, the start of the campaign is no longer relevant, as long as it happens within a 90-day window

Interestingly, our results showed that, once 22 days have passed after the publication of the article, the time-lapse size to start the campaign was not important for reaching a relevant number of mentions in the news. This delay had limitations in the case of PR articles, since *Eurekalert!* only allowed posting news about publications that are maximum 90-day old.

Then, independently of the timing, X posts and mentions in the news (and therefore, AAS) always increased immediately after the communication campaign, no matter how much it was delayed within the time window of the experiment (**Fig. 13**).

**Fig. 13.**
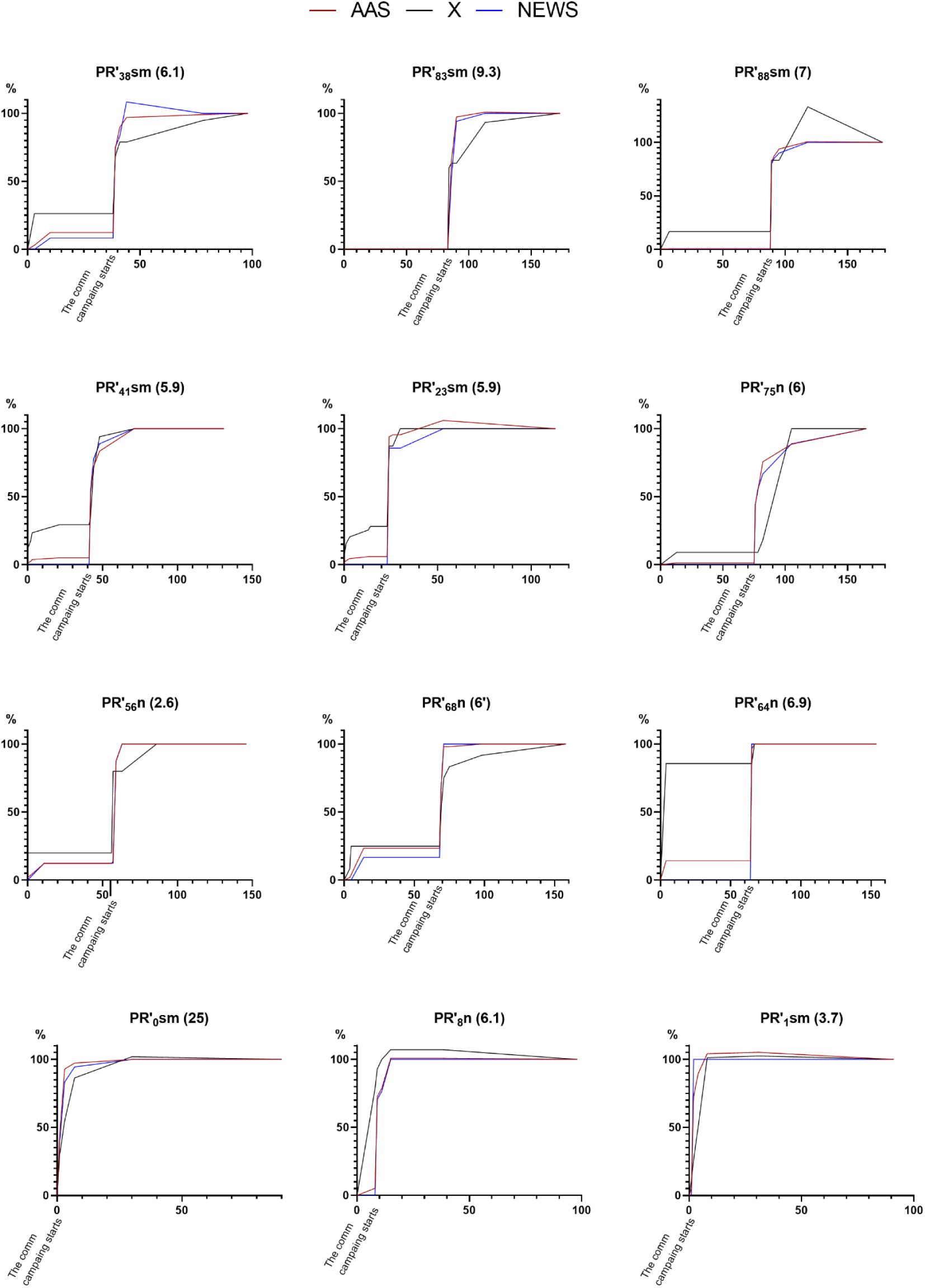
X posts and mentions in the news increase independently of the campaign timing. % of increase in the AAS, number of X posts (according to Altmetric) and mentions in the news (according to Altmetric) since the publication day, for PR’ articles (PR’_+22_ and PR’_-22_). The numbers in parentheses refer to the impact factor of each article. *Note: News mentions are shown in their raw form, including false positives from Alzforum, EurekAlert!, and AlphaGalileo, since these mentions are counted in the AAS and omitting them would make the interpretation of the observed AAS confusing*.

Finally, we also examined the potential effect of additional dissemination actions carried out months after the initial campaign, through summer reminder posts on X (one post per article in August 2022).

Interestingly, the effect of these one-off actions was minimal, and the AAS of these articles increased by no more than 1 point one week later (**Table 11**).

**Table 11.**
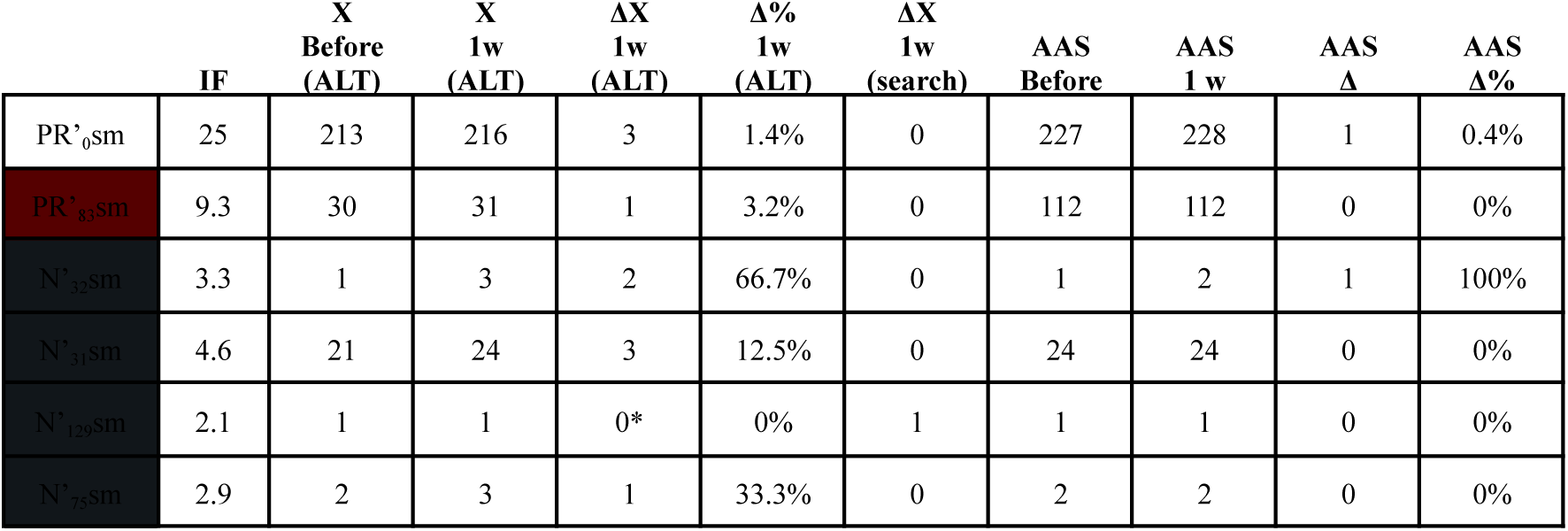
Evolution of the X mentions and AAS of each article after a summer reminder on X. The subscript in the reference for each article indicates the number of days between the article’s publication and the start of the advertising campaign. *Note: As happened in our first campaign, Altmetric did not detect one of our tweets, although it was correctly referenced (it is marked with an asterisk and counted in the search column)*.

#### 3.3.5 One week after the campaign, PR’_+22_ articles increased their abstract reads by 10 times in contrast with NPR’_+22_ articles

Articles in the PR’_+22_ group experienced an average increase of 482 reads per article (a 72% increase) one week following the campaign, with the PR’_+22_sm subset showing the highest increase (**Table 12**). In contrast, articles in the NPR’_+22_ group only saw an average increase of 51 reads per article (a 12% increase). However, it is important to mention that not all articles could be analyzed, as only some editorials provided the necessary data.

**Table 12.**
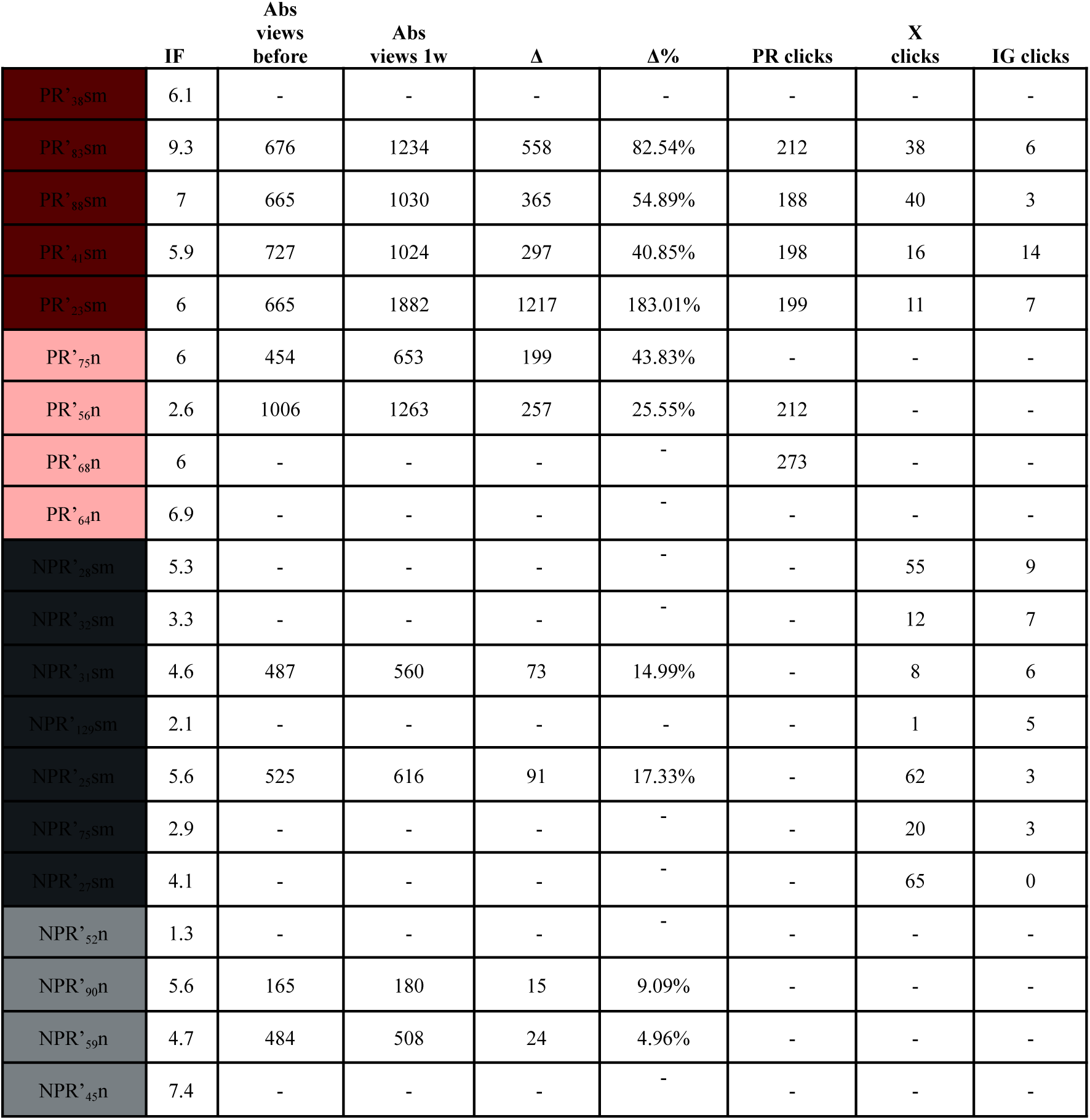
Number of reads and clicks one week after the communication campaign (only for those articles for which the information is available). The subscript in the reference for each article indicates the number of days between the article’s publication and the start of the advertising campaign. *Note: The article PR’68n corresponds to article n(6’) in graphs 10c, 11c, 12d, and 13*.

To determine the effects of the PR or SM on this increase, we conducted an inter-subject analysis and found that PR was the only significant factor influencing the number of views [F(1,5)=12.357, p<0.05]. All other factors and interactions, including IF and SM, were not significant. Also, the decomposition of the interactions reveals that, prior to the intervention, the differences between the groups were not statistically significant.

We also identified where the number of clicks came from (either from the PR, X posts, or IG posts), and found that PR links were the most prevalent, clicked by an average of 214 times compared with X links and IG links, that were only clicked 30 times, and 6 times respectively (**Table 12**). These results, once more, indicate the relevance of delivering a PR digesting the scientific content of a neuroscience article to increase its social attention and influence.

Overall, our results highlight the importance of the dissemination campaign in the field of neuroscience, and especially the generation of specific materials to be released to the press to increase social attention.

## 4. DISCUSSION

### 4.1 Sending a press release is the most effective way to be mentioned in the news and, therefore, increase the AAS of an article

This study supports that the optimal communication strategy to attract media attention to a scientific article is by elaborating and issuing a PR summarizing its content. When we retrospectively analyzed the 2016–2021 INc-UAB articles, we observed that those accompanied by a PR (PR group) received 39.5 times more mentions in the news, which was reflected in a 71-point difference in their AAS compared to those without a PR (NPR group). Most importantly, the same pattern was observed in the prospective study. Articles with a PR (PR’ group) were the only ones that received relevant media attention, with an average of 18.6 mentions in the news per article (counting those detected and undetected by Altmetric). These mentions appeared in 99% of cases after the PR was issued, even though there was at least a 23-day gap between the publication date and the release, demonstrating the key role of the PR on the media attention. PRs also led to an increase in X posts and article reads, as well as a 68-point growth in the AAS. The fact that both analyses show a similar AAS difference indicates the consistency in the impact of the dissemination campaign and the accuracy of the methodology employed.

These results fill an important gap highlighted in a recent publication, in which Fuoco and colleagues (2023) performed a retrospective study concluding, as we do, that the primary factor influencing online attention is the issuance of a PR. However, in their article, due to the type of analysis performed, they noted that causality could not be ensured. In this sense, since we developed a prospective study comparing articles before and after different communication strategies, we could look at the direct effect of each campaign, demonstrating that PRs are key elements to expanding social attention. This was true even for high-IF articles: although they might initially appear to be of interest to the general public, they received no attention if a PR was not issued.

An important point, however, is that not all articles with a PR received the same level of media coverage, even though we could consider all of them potentially newsworthy. The extent of coverage and the subsequent increase in their AAS (around 70 points on average in our case) likely depend on multiple factors, such as the research field (i.e. Neuroscience), the type of institution issuing the PR, and other contextual elements. Additionally, the way the PR is crafted (i.e. structure, type of language, etc.) may significantly influence its effectiveness, which is particularly interesting and would need further investigation in future studies.

### 4.2 If the press release is not issued before or immediately after the publication of the article, the delay does not seem to affect social attention

Although PRs are usually issued around the time of publication, our findings suggest that even when there is a delay, significant social attention can still be achieved. This complements what Fuoco and colleagues previously described (Fuoco et al., 2023), showing that, aside from the release of a PR, other variables played an important role in media coverage, such as the day of the week the article was published, the type of access (pay-per-view or open access), and how promptly the PR was issued. According to them, PRs have more media attention when they are delivered on the same day of publication, and decay when they happen more than one day after. Our results suggest that after this immediate time window, it does not matter how much a press office waits to start the communication campaign, always keeping in mind that *Eurekalert!*, one of the main platforms to post PRs, only allows material from 90-day old articles maximum (Eurekalert!, 2018).

In some cases, it may be worthwhile to wait a few weeks until there is a special occasion, as it might result in a higher impact. For example, if the article is about Alzheimer’s disease, it would be smart to wait until Alzheimer’s disease day, if possible. According to our study, selecting the best moment to distribute the PR would not be a problem, within a reasonable time window.

Once the PR is distributed, as we observed in our prospective experiment, mentions appear within the next hours or days (both in the news and on SM). This provides further clarity to the findings of Fang & Costas (2020), who demonstrated that the average news half-life (which refers to the number of days until 50% of all mentions have appeared) is 22 days. With the accumulation patterns that we observed, we suggest that, if this is the case, it is not because it takes an average of 22 days for journalists working in media outlets to prepare a piece of news, but because it must probably be the average number of days it takes for press offices to send the PR.

### 4.3 Single X posts have limited influence boosting online mentions

Our retrospective study of in-house articles (study 2) showed that for articles without a PR (NPR group) there was a significant but modest association between X posts, media coverage, and increased Mendeley readership (but not citations), suggesting that the last two might be driven by SM exposure. However, in the prospective study we found that isolated X actions had no effect on online attention. ANOVA and ANCOVA tests indicated that single posts did not lead to significant differences in the counts of news items about the article, X posts, article reads, or the AAS; instead, these changes were attributable to the PR.

Concerning the AAS, it is important to note that, due to the Altmetric weighting system (Altmetric, 2023), a substantial increase in the number of X posts would be needed to observe significant differences. However, as just mentioned, the X interventions in our study not only caused no significant differences in the AAS, but also did not affect the number of X posts, indicating that the weighting system is not the main reason for the lack of effect observed here. Future studies with larger sample sizes, more intensive interventions, and a stricter X-posting schedule may provide better insights into the effects of posting on X.

Moreover, a deeper analysis that also considers post engagement would be of great interest. As Fang et al. (2022) and Martin and MacDonald (2020) noted, user interactions with posts mentioning scientific articles are generally low. In this context, a high number of tweets does not necessarily ensure that the information reaches the intended audience. Instead, measuring how users engage with these tweets (through retweets, likes, replies, or link clicks) would capture SM attention in finer detail (Fang et al., 2022). Composite altmetric tools, such as Altmetric, should consider this, as it does not appear to be explicitly included in their weighting scheme when calculating scores (Altmetric, 2023, 2024).

Regarding why SM posts often fail to generate engagement with the public, the reasons are likely multifaceted and complex. A key factor must be the generally low quality of the text of most tweets, as many contain only the article title or a formulaic text devoid of originality (Eysenbach, 2011; Robinson-Garcia et al., 2017). This is neither engaging nor easily understandable for a general audience, and suggests that, rather than communicating their research to society, researchers are primarily using X to publicize their work among peers (Tsou et al., 2015). If alternative metrics aim to capture an article’s impact beyond academia, such tweets should ideally be filtered out, as suggested by Robinson-Garcia and colleagues (2017).

### 4.4 Further research is needed to understand the dynamics between social media, readership, and citations

Regarding the factors influencing the number of abstract reads, the ANOVA tests suggested that while PR may have an effect, this influence was not observed for X isolated posts. Despite the limited data in our study, these results are consistent with previous research on SM.

As briefly mentioned in the introduction, Fox and colleagues (2015, 2016) found no significant variations in the number of reads or downloads when comparing articles from the *Cardiology* journal (IF=15.2) posted on the journal’s SM to those that were not. Similarly, Tonia et al. (2016) analyzed the effects of disseminating articles from the *International Journal of Public Health* (IF=2.3) via the journal’s blog, X, and Facebook, and found no changes in the number of downloads or citations. More recently, Branch et al. (2024) also found no citation increase linked to promotion on X, although tweeted articles did see a significant rise in downloads.

Our first retrospective analysis does reveal a correlation between tweets and citation counts. However, it remains unclear whether this relationship reflects a causal influence (where SM increases visibility and, therefore, citations) or merely indicates that articles of greater societal relevance are also more likely to be cited. Long-term prospective studies are needed to determine the true impact of communication campaigns on citation outcomes, ideally also considering the engagement with the tweets. Retrospective analyses similar to our second study, but with larger sample sizes, could also yield valuable insights. For instance, by comparing whether a high AAS resulting from PR correlates with citations as strongly as a high AAS driven by active discussions on X, it would be possible to determine whether scientific conversations on X remain confined to researcher-only bubbles (whose members may later cite these articles in their publications) or extend to a broader audience.

With regard to the factors that may explain why X activity alone has limited influence on citation counts, several reasons may be involved, stemming from both platform design and user behavior. Firstly, X algorithms prioritize trending or broadly appealing content over scholarly relevance, reducing visibility among academic audiences. Secondly, posting a tweet mentioning an article does not ensure that either the user posting it or the audience will access or read it. Finally, researchers may rely on other mechanisms to stay up to date with new publications, such as journal alerts, newsletters, or academic databases, meaning that social media promotion may not significantly affect their awareness of a given article.

Nevertheless, as Branch et al. (2024) argue, even if X does not directly increase citations, it can still play a valuable role in researchers’ professional lives by fostering collaboration, enabling knowledge exchange, and opening up career opportunities through the cultivation of online academic communities. Further studies should explore these broader benefits in more depth.

Finally, we would also like to mention that due to the controversial political decisions associated with Elon Musk (the current owner of X), many individuals and media outlets have recently abandoned the platform. Alternative social networks are emerging, such as Bluesky and others. Particularly, Bluesky operates in a very similar manner to X, leading us to believe that the conclusions drawn in this study would also apply to this new platform, as well as to other similar networks that may arise in the future.

### 4.5 Artificial intelligence tools may be useful in the future when preparing dissemination materials, although some aspects need to be considered

Preparing dissemination materials requires time and dedication. For a PR, we calculated an average of 9 hours, and 2 hours for SM posts (less for those with a PR, as ideas can be reused). This may change with artificial intelligence tools, which in the future may be a great instrument to generate first drafts of PRs, or SM posts in a more agile way. Nevertheless, beyond the existing technical limitations of AI tools, challenges pertaining to copyright, sources trustability, and the underlying training data utilized by these systems still require resolution. In fact, *Eurekalert!* updated in January 2024 its Content Eligibility Guidelines, to expose that, due to all these reasons, they will not accept AI-generated text, images, and other multimedia for submission and distribution via the platform (Eurekalert!, 2024). However, even if these concerns are finally settled, others may appear: AI tools helping to generate thousands of PRs a day could limit their efficacy by saturating journalists and ultimately the users.

### Recommendations for researchers and press offices

Based on our findings and the current context, we propose the following recommendations:

● **Prioritize PRs:** Sending a PR is the most effective way to increase media attention. PRs consistently outperform SM efforts in driving public engagement. However, not all articles are inherently newsworthy or suitable for PRs. Future research could explore the factors that determine newsworthiness to better identify articles of general interest.
● **Timing matters, but flexibility exists:** While issuing a PR on the publication day maximizes visibility according to existing literature, a delay beyond this initial window may not significantly impact media attention. Strategic timing (such as aligning with relevant awareness days) may enhance impact.
● **Single SM actions are insufficient:** Individual posts on platforms like X have a limited impact on overall online attention. While SM can enhance researchers visibility, facilitating networking and collaborations, and may even weakly influence citations over time (a relationship that still requires further investigation), direct journalist outreach through PRs remains the primary driver of public and media interest.
● **Ensure that digital mentions include proper references:** To allow altmetric tools to accurately track mentions, researchers and institutions should include a direct citation of the article when sharing it on SM, institutional websites, or other dissemination platforms. Failing to do so may result in lost tracking opportunities and an underestimation of the article’s true reach.

### Altmetric limitations

Although AAS represents a powerful tool to measure the social attention of a scientific article, it contains some limitations. On the one hand, due to its definition and characteristics, it ignores mentions in offline media, such as television, radio, or print newspapers. Also, because it is constructed as a single composite score, it could mask relevant differences between cases (i.e. two papers may end up with the same AAS, yet that number could stem from very different patterns of attention).

On the other hand, as we observed in our study, Altmetric failed to detect at least half of our articles’ mentions in digital news. Fleerackers and colleagues (2022) had more accurate results, finding that Altmetric recalled 71% of them. However, as in our study, a larger proportion was missed. As declared in the Altmetric website, they do not automatically capture mentions across the entire web but rely on a “manually curated collection of sources” (Altmetric, 2020), omitting part of the coverage. This, together with the methods that Altmetric uses to identify mentions: a mixture of link matching and text mining (Fleerackers et al., 2022), may account for the loss of data.

News stories often lack a proper citation of the article (authors, title, or DOI), which complicates the detection for Altmetric. Fleerackers and co-workers (2022) analyzed 8 news sources and showed that only 57% of their stories properly cited the research study (namely, link to the publication, institution, authors, journal, etc.), whereas we found in the present study that it was merely 45%. If journalists included the article reference in their pieces, it would not only improve Altmetric precision but it also would benefit readers, being able to reach the original article easily.

Plus, the language used in the pieces mentioning the article (both in news outlets or in SM) can be a barrier for Altmetric detection. According to Ortega (2020, 2021), Altmetric has a preference for news outlets published in English as well as for topics of general interest. Since many of the pieces of news and posts referencing our articles were written in Catalan and Spanish, this could have potentially reduced its effectiveness. In fact, there was a mix of reasons making it more difficult for Altmetric to identify pieces in other languages in our study: these were also those that most often failed to properly cite the publication. Altmetric detected 97% of the news items in English, including those without a reference to the article, but identified only 10% of the pieces in other languages, failing to detect 97% of those without a reference.

We also detected that Altmetric may contain some misleading counts in its score. For example, we observed that they considered “mentioned by news outlets” articles appearing on online lists of papers (i.e those listed in https://www.alzforum.org/papers) or PRs posted online by communication departments on *Eurekalert!* and *Alphagalileo*, which could be regarded as noise and should be filtered out.

Finally, with respect to X, posts sharing a piece of news about an article, but without proper reference to the publication itself, are not taken into account by Altmetric. This means, for publications with a PR, that around 75% of the conversation is neglected in the AAS. Researchers and institutions should be aware of this, and ensure that their posts include proper citations. Additionally, three of our posts were not detected either, even though they correctly referenced the article.

Despite all these drawbacks, Altmetric data collection on news is perceived as less biased compared to its competitors, such as PlumX (Ortega, 2020, 2021). According to Ortega, Altmetric has a more geographically and linguistically diverse coverage, whereas PlumX is more concentrated in the United States. Besides, Altmetric includes a greater variety of general-interest news outlets, while PlumX tends to emphasize a narrower selection of sources.

Concerning the general limitations that alternative metrics may have, some researchers have warned of the risk of unethical manipulation. They can be improved by self-citations on SM, sending a PR, etc. (Scarlat et al., 2015), which may not be reflecting a genuine social interest but the amount of effort to build an efficient communication campaign. In our opinion, this does not have to be perceived as a problem: disseminating research results is one of the duties that at least researchers working in the public system have, and altmetrics reflect numerically the campaign success. However, these particularities have to be taken into account when defining the use and interpretation of alternative metrics tools.

We envision that in the future, new open indexes to evaluate research, promoted by public research entities, will probably be developed, and they will have to consider the importance of including the social interest. We are certain that AAS advantages and limitations surfaced in this article may be of interest to improve and generate novel tools to measure scientific and social attention.

### Study limitations

Our prospective experiment has certain limitations. First, as the study focuses on real communication campaigns at a relatively small research center, we had access to a modest sample size. However, we consider that the consistency of the results mitigates concerns about scale. Second, although the articles vary in terms of topic, study type, journal, and access format, all are within the neuroscience field, ensuring a relevant degree of homogeneity. Lastly, we analyzed articles published and PRs issued at different times throughout the year (as shown in Table 1). Although this variability is not ideal, as suggested in previous studies (Fuoco et al., 2023), such factors do not undermine our key conclusion: PRs are instrumental in maximizing the societal impact of scientific research.

## Author contributions

All authors contributed to the study conception and design. Data collection was performed by J. Giner-Miguelez and R. Bastida-Barau. Data analysis was carried out by R. Bastida-Barau. The first draft of the manuscript was written by J. Giner-Miguelez and R. Bastida-Barau, and S. Ferré and C. Barcia contributed to the final writing and edition of the article. All authors read and commented on updated versions, and approved the final manuscript.

## Acknowledgements

We would like to thank professor Roser Nadal (INc-UAB) for her invaluable assistance with the statistics, the Quintanalab (INc-UAB) for their priceless advice, and Maria Jesús Delgado and Octavi López (UAB Press Office) for their efforts in compiling past press releases and their continuous help. We are also deeply appreciative of the support provided by Simon Bilevych and Mariam Lepage (Altmetric), who have been on the other side of numerous emails addressing our questions about the platform with great kindness and efficiency. We finally extend our heartfelt gratitude to Brian Lin and Ashley Phan (EurekAlert!), Lizany Thomas and Eilidh Ruthven (Wiley), Bhavani Mutharasan (Elsevier), Facundo Santome (MDPI), and Hadley McIntosh Marcek (ACS Chemistry for Life), whose invaluable assistance in gathering essential data for this study was marked by courtesy and dedication.

